# The accessible human surfaceome

**DOI:** 10.64898/2026.10.18.759859

**Authors:** Rebecca J. Carlson, L. Robert Hollingsworth, Michael J. Smallegan

## Abstract

Proteins associated with the extracellular face of the plasma membrane (“surface proteins” or the “surfaceome”) are important therapeutic targets because they can direct therapeutics to specific cell types and deliver them into cells. However, experimental and computational databases disagree substantially on which proteins make up the human plasma membrane proteome, and none focus specifically on the set of accessible surface proteins that would be useful for therapeutic applications. Here, we used an agentic LLM pipeline to curate the accessible human surfaceome from literature. To assess performance, we built SurfaceBench, a manually curated benchmark of 147 proteins on which five reference databases frequently disagree. Our LLM triage agent classified surface status more accurately than existing databases and recovered 1,108 candidate surface proteins not present in any of them. On held-out clinical and viral-entry surface targets, the pipeline recovered 98–100% of established surface antigens. A literature-anchored deep-dive pipeline run on over 5,000 candidates then returned 1,783 canonical surface proteins and 1,252 with weaker or context-dependent surface evidence. Each queryable record contains 24 LLM-based annotation fields — including confidence, primary localization, and expression breadth — and seven deterministic structural and topological features. Unlike existing databases, our records trace every claim to the agent’s reasoning and to exact quotes from the full-text literature it could access. All records can be browsed in an opensource viewer. These results show that LLMs reasoning over unstructured scientific text can build updatable, bespoke databases that outperform expert-curated resources, although performance remains limited by agent access to full-text literature.

## INTRODUCTION

Surface proteins have important biological functions, since their accessible location allows cells to communicate with each other and with their environment. Only an estimated 10–20% (“Plasma membrane (HPA),” 2026; Thul et al., 2017) of the human proteome is at the cell surface, but more than 65% of approved drugs (Laurent et al., 2025) target these proteins. Yet “surface protein” is difficult to define. Some proteins that are typically intracellular reach the surface only in specific cell states: the ER chaperone BiP (GRP78) relocates to the surface under ER stress (Zhang et al., 2010), and the lysosomal membrane protein LAMP1 appears on cytotoxic T cells only as they degranulate (Betts et al., 2003). Other proteins are surface-exposed on a single cell type, like the sperm fusion protein IZUMO1 (Inoue et al., 2005), or are secreted but held at the surface by a membrane partner, as MD-2 is by TLR4 (Loes et al., 2021). Whether a protein counts as surface resident therefore depends on the context a scientist cares about.

We define the accessible surfaceome as the set of human proteins that can be reached from the extracellular face of the plasma membrane in at least one experimentally validated context. This includes proteins accessible only in specific cell types or states, which are often the most biologically and therapeutically interesting. We focus on accessibility rather than membrane localization because it determines whether anything outside the cell (a ligand, a neighboring cell, or a therapeutic) can engage the protein.

Existing surface protein databases were not built around this definition. UniProt (UniProt Consortium, 2025) and Gene Ontology Cellular Component (GO CC) (Gene Ontology Consortium, 2026) are curated, whereas the Human Protein Atlas (HPA) (Thul et al., 2017) and the Cell Surface Protein Atlas (CSPA) (Bausch-Fluck et al., 2015) are experimental. SURFY (Bausch-Fluck et al., 2018) is predictive and not fully independent, as it was trained on transmembrane proteins supported by CSPA, UniProt, and the COMPARTMENTS database (see Methods). HPA’s antibody atlas spans both membrane leaflets, while CSPA detects only glycosylated outer-surface proteins. SURFY’s definition is closest to ours, but it scores only predicted transmembrane and select GPI-anchored proteins from sequence features, so it can miss noncanonical surface presentation documented in the literature. As a result, the five databases agree on only 188 surface proteins out of a union of 5,546 proteins classified as surface in at least one database (Figure 1). Each also returns a fixed, steady-state call without its supporting literature, so users must still search the literature to decide whether a listed protein suits their question.

**Figure 1.**
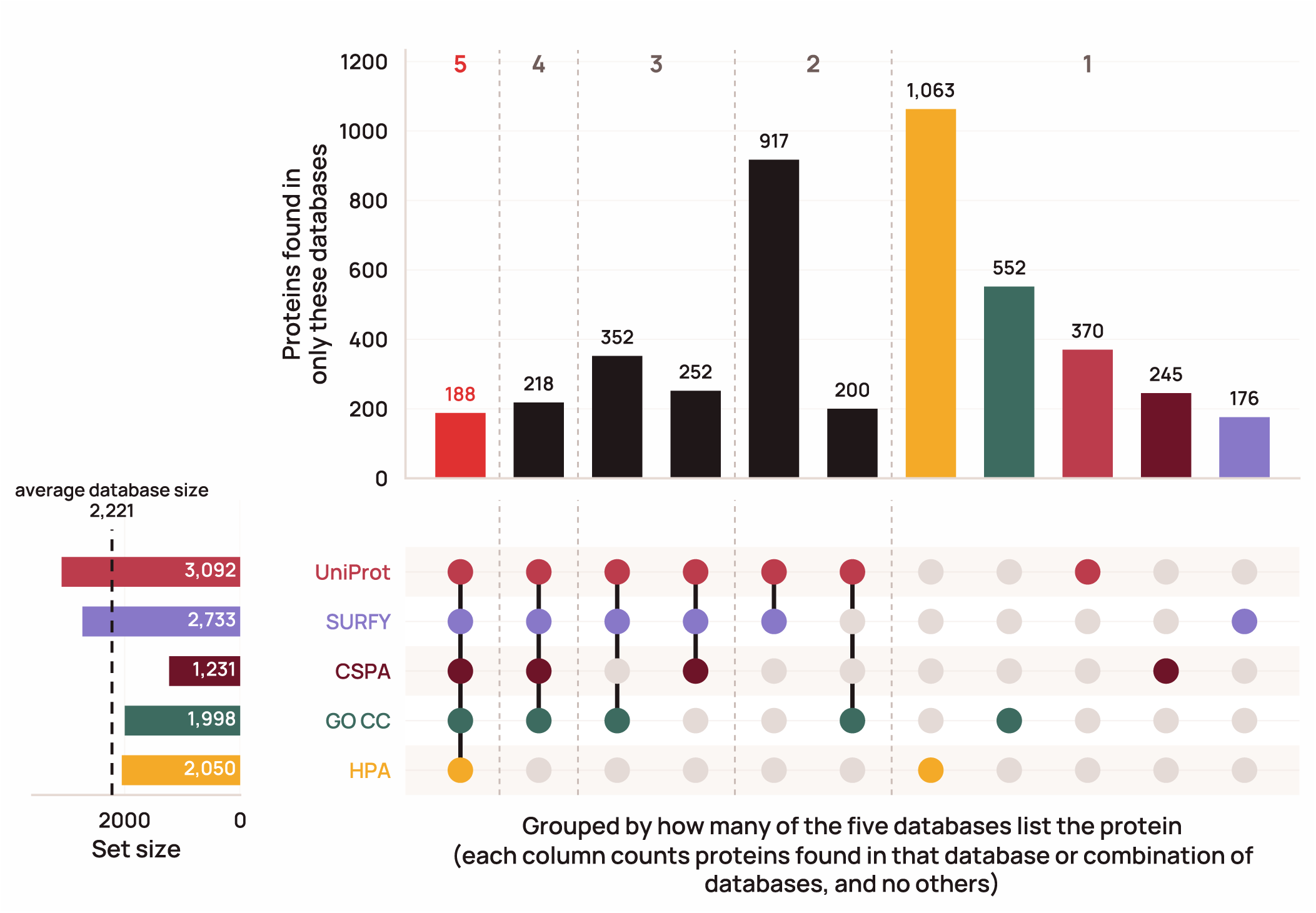
Five surface protein databases agree on only 188 proteins. UpSet plot of five surface protein databases; each bar counts proteins found in that database or combination and no others, grouped by how many databases list the protein, with total set sizes shown on the left. Bars are shown for the 10 largest combinations plus each database alone. Each set comprises the protein-coding genes with at least one UniProt accession labeled as a surface protein under the source-specific rule (UniProt: strict surface terms or extracellular topology; GO CC: at least one non-electronic (non-IEA) annotation to a surface-relevant term; HPA: Enhanced/Supported/Approved reliability for Plasma membrane or Cell Junctions; SURFY: published binary classifier output; CSPA: high-confidence or putative; details in Methods). [Figure data and script](https://gist.github.com/beccajcarlson/d655abfc9c7deeaff1cfbe584de96ffa).

Previous approaches for assembling literature context are limited in scale and quality. Expert curation cannot keep up with the literature, which grows faster every year. Structured search and text mining can find proteins that are mentioned together with surface terms, but neither can judge the context of a claim. Meanwhile, LLMs can now read full-text papers at proteome scale, decide whether a claim supports surface accessibility and under what conditions, and enable tracing of the reasoning and verbatim evidence behind each verdict.

Here, we used an agentic LLM pipeline to curate the accessible human surfaceome from the literature. To ensure coverage, we first ran a lightweight triage agent, with a brief prompt and no tool calls, on the whole human proteome (19,324 protein-coding genes), classifying each as surface, contextually surface, or non-surface with a structured reason for every call. On SurfaceBench, a manually curated 147-protein benchmark enriched for database disagreement, it classified surface status more accurately than any of the five reference databases, individually or in combination. Then, for proteins classified as contextually surface or surface by our triage agent or classified as a surface protein in at least one of the five reference databases, we ran a deep-dive pipeline. This pipeline instructed agents to read the retrieved literature and return a structured record that anchors each claim to verbatim quotes. This yielded 1,783 “canonical” surface proteins and 1,252 likely surface proteins, each with 24 queryable LLM-derived annotations and deterministic structural and topology features.

We did not impose a single definition of the surfaceome, because each verdict comes with its reasoning. Alongside the overall surface call, each record reports evidence strength, cell-type restriction, state dependence, and extracellular domain size as separate fields that can be queried directly. Our canonical and likely tiers are a starting point, but users can also assemble the surfaceome that fits their own question, for example by selecting proteins that are constitutively exposed across most tissues or understudied proteins that may represent new surface biology. All records, code, and data are archived on Zenodo and available through an open-source viewer at surfaceome.deliverome.org and a public API (Methods).

## RESULTS

### A triage agent classifies hard-to-categorize surface proteins more accurately than existing databases

We first asked whether an LLM could triage the whole proteome from its own knowledge, leaving deep literature grounding for a second, evidence-anchored pass. To test this, we created a 147-protein benchmark ([SurfaceBench](https://surfaceome.deliverome.org/benchmark/)) enriched for database disagreement on surface protein status. We manually labeled each protein as surface, contextually surface, or non-surface, with a reason from a 19-category taxonomy (Table S1); for example, a protein may be contextually surface because it is cell-type restricted or cell-state induced, or classified as non-surface because it is endomembrane-resident or nuclear. The benchmark set was biased by design toward database disagreement to enable detection of performance differences within the constraints of manual benchmark annotation, so we also later evaluated proteome-wide calls on held-out sets of clinical and viral-entry targets (see below). Each database defines surface membership differently, so we re-derived inclusion rules for each database against SurfaceBench and kept the most accurate variant for each to enable a fair comparison (Figure S1).

We then defined a brief [triage prompt](https://surfaceome.deliverome.org/prompts/#triage-system) (Methods) that asks whether the protein is accessible from the extracellular face of the plasma membrane. The response requires a verdict, a reason from the taxonomy, a confidence level, and a brief rationale. Because a non-surface call can remove a protein from all downstream analysis, before calling a protein non-surface the model was required to name each potential contextual reason and cite the protein-specific evidence that rules it out.

We compared Claude models on SurfaceBench performance. Given only the gene symbol, Sonnet 4.6 classified 95.7% of SurfaceBench proteins correctly, nearly matching Opus 4.8 (95.9%), and well ahead of the smaller Haiku 4.5 (81.6%) (Figure S2). Because triage runs on every protein-coding gene, we chose Sonnet 4.6, the cheapest model at this accuracy plateau (Figure S3). Pre-resolving each gene to its NCBI Gene, HGNC, and UniProt identifiers improved accuracy on every model: modestly on Sonnet 4.6 (+1.8%) and Opus 4.8 (+2.3%), and substantially on Haiku 4.5 (+8.6%) (Figure S2). Identifiers primarily improved recall of “yes” and “contextual” calls (Figure S4). Adding a web-search tool or retrieved PubMed abstracts did not significantly increase accuracy (exact McNemar test, p ≥ 0.34) but substantially increased cost (Figures S2, S3). Sonnet 5 with identifiers performed slightly, though not significantly (p = 0.29), worse than Sonnet 4.6 at higher cost (Figures S2, S3), so we did not use it. Most of this work was completed before the Claude 5.5 models were released; when we re-ran the benchmark on them with identifiers, Haiku 5.5 and Sonnet 5.5 both matched or exceeded their predecessors at equal or lower cost, while Opus 5.5 declined to answer about 5% of calls, all on surface proteins (Figures S2, S3).

Compared to the five reference databases, Sonnet 4.6 with no external tools classified surface status more accurately than any, whether individually or as an ensemble (Figure 2; Figure S5). Notably, UniProt with an optimized cutoff performed very well, especially on clear yes or no cases, achieving 90% overall accuracy. Unlike the other sources, each of which relies on one class of information (experimental evidence, curation, or a sequence model), UniProt called a protein surface if curators placed it at the surface or if it had a transmembrane domain or a signal peptide, and incorporating multiple signals with the benchmark-tuned cutoff increased its performance across diverse proteins.

**Figure 2.**
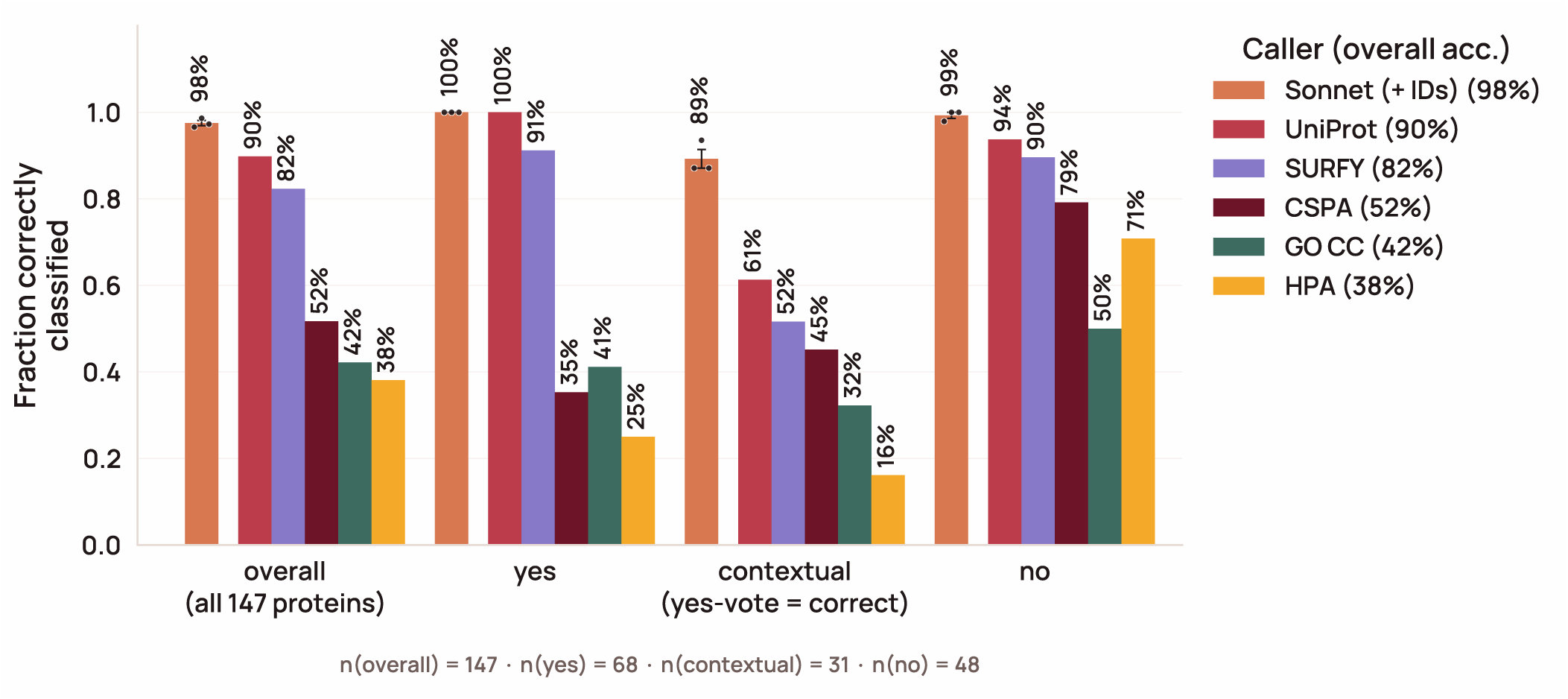
A tool-free accessibility triage agent classifies hard-to-categorize surface proteins more accurately than any single database. Classification accuracy on SurfaceBench, 147 proteins enriched for disagreement among the five source databases and manually labeled as surface, contextually surface, or non-surface. Bars compare the triage agent (Sonnet 4.6, no external tools) against the five source databases; contextually surface proteins are counted as correct when called surface. Sonnet 4.6 was run with three replicates. Error bars indicate SEM. [Figure data and script](https://gist.github.com/beccajcarlson/2bb4f7aac629535982c142bc2032e04d).

Sonnet matched or exceeded optimized UniProt within each of the three verdict categories (Figure 2), so its advantage does not come from a permissive bias toward surface calls. In addition to verdict agreement, the agent’s specific reasons for the surface call (e.g., endomembrane vs nuclear within the “no” category) also usually agreed with the curator’s, with an overall reason-specific accuracy of 88% (Figure S4). The largest single-category disagreement (4 of 18 errors) was cytoplasmic kinases (AKT2, BTK, JAK1/2) that the agent called “inner-leaflet anchored” rather than “cytoplasmic.” These kinases indeed fall between the two provided categories (the JAKs, for example, sit constitutively on cytokine receptor tails but carry no anchor of their own), and all four remain labeled with the correct overall “no” verdict.

### Proteome-wide triage recovers surface candidates missed by all five databases

We next ran the triage agent across the full set of protein-coding genes (19,324 genes) in two stages. First, the tool-free agent (Sonnet 4.6 with identifiers) classified 2,528 as surface, 1,721 contextually surface, and 15,075 non-surface at a total cost of $116 ($0.006 per gene). To maximize sensitivity, we re-triaged every “non-surface” call supported by no database and every high-confidence “non-surface” call supported by exactly one database (n = 14,404), this time with relevant PubMed abstracts supplied in the prompt so that a documented surface proteoform could be identified from a high-level literature survey. KLK2 was one example of a candidate surface protein rescued through this approach: it was long considered secreted, but a membrane-bound form is enriched in prostate cancer and is now targeted in Phase 3 clinical trials (Chi et al., 2026). The tool-free agent called it non-surface; with PubMed abstracts, the agent reclassified it as contextually surface. Across the proteome, literature volume had little effect on these second-stage reclassifications, with rates across quintiles of gene2pubmed-linked publications ranging from 1.6 to 3.3% without a consistent trend.

In total, the two-stage triage recovered 1,108 candidate surface proteins absent from all five databases, with 314 contributed by the PubMed stage. Only 87 of these were called definitively surface, while the rest were contextual (Figure 3), indicating that, as expected, the databases mainly miss state- and lineage-dependent surface biology rather than classical surface proteins. Examples of rescued surface proteins include LY96/ MD-2, a secreted protein that stably associates with TLR4 as a co-receptor on the cell surface (Loes et al., 2021), the surface pore-forming protein GSDME (Wang et al., 2017), the outer-leaflet ganglioside sialidase NEU3, which has no predicted transmembrane helix (Rodriguez-Walker and Daniotti, 2017), and NPM1, a nucleolar chaperone with a surface pool reported on AML blasts that is targeted by a preclinical antibody (George et al., 2026a) (Figure 3).

**Figure 3.**
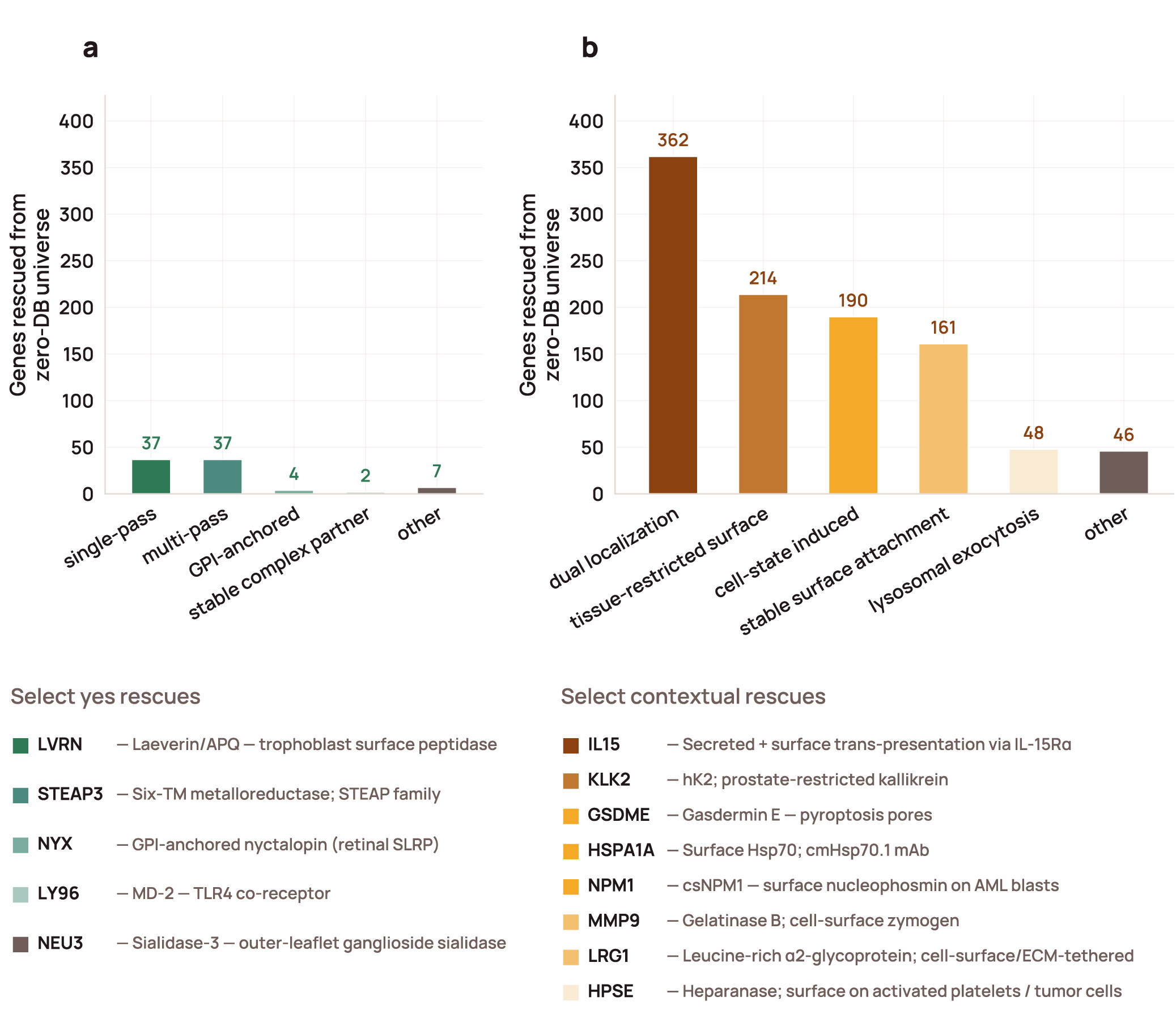
The triage agent rescues 1,108 surface candidates not identified by any of the five source databases. The two-stage Sonnet 4.6 triage agent calls 87 proteins definitively surface (“yes,” panel a) and 1,021 proteins contextually surface (“contextual”: state- or lineage-dependent, panel b) that were not identified by any of the five source databases. Bars group each verdict by the agent’s rationale, with representative examples labeled. [Figure data and script](https://gist.github.com/beccajcarlson/a4526c9e6de5e958826bf1d764744c1b).

Because no hand-curated ground truth exists at proteome scale, we scored the reference databases against the agent’s proteome-wide calls as a reference (Figure S6). Agreement ranged from 75% (HPA) to 90% (UniProt), with UniProt agreeing more closely than any combination of databases. Many disagreements reflect our accessibility definition itself. Among database-annotated proteins that the triage agent declined, the leading reasons were cytoplasmic localization (42%), inner-leaflet anchoring (20%), and endomembrane residence (19%): the databases largely capture proteins associated with the plasma membrane rather than those that are extracellularly accessible. For example, 73 proteins received surface calls from three or more databases but a triage “no”, including membrane-associated scaffolding proteins that bind the plasma membrane from the cytoplasmic side, such as EZR, MSN, and DLG1.

To check these proteome-wide calls against standards independent of SurfaceBench, we scored the two-stage triage verdicts on three orthogonal target lists expected to be mostly surface proteins: 1) curated preclinical and clinical antibody–drug conjugate (ADC) targets from three existing databases (Buniello et al., 2025; Raybould et al., 2020; Shen et al., 2023), 2) preclinical and clinical T-cell engager (TCE) targets (Raybould et al., 2020), and 3) viral-entry receptors from ViralZone (Hulo et al., 2011) (see Methods). Sonnet recovered more of these targets than any reference surface protein database: 98–100% of each set (324 of 326 overall), compared with 92–97% for UniProt, still the best single database (Figure S7). The only two targets the agent did not call surface, both of which we consider correct non-surface calls, were ADC target CT83, a poorly studied intracellular cancer-testis antigen named only in a single ADC patent and not firmly established as a surface protein (patent #WO2026142343), and viral receptor NPC1, a lysosomal receptor with little established surface localization that assists with Ebola virus lysosomal escape rather than initial uptake via macropinocytosis (Carette et al., 2011).

These target sets contain only established surface proteins, so they test recall but not specificity. To ask whether SurfaceBench accuracy should transfer proteome-wide, we compared the benchmark’s topological composition with that of the agent’s proteome-wide surface calls (Figure S8). GPI-anchored and glycosylated proteins were over-represented in SurfaceBench, while multi-pass transmembrane proteins and those lacking a signal peptide were under-represented, but no broad category was absent. Because the benchmark is weighted away from the easier-to-classify multi-pass proteins, its accuracy is likely a conservative estimate.

We then used the same topology features to compare surface calls from Sonnet, from each database, and from the Sonnet-only set with and without the PubMed stage (Figure S9). Database calls were dominated by proteins with classical membrane-topology signatures (transmembrane helices, GPI anchors, and N-glycosylation sites), and the mass-spectrometry-based CSPA was the most biased toward glycosylated proteins. Sonnet-only calls were depleted of all three, so the agent mainly contributes surface proteins that lack canonical topological features, which prediction-based and experimental databases miss.

### A literature-anchored deep dive produces evidence-backed records for candidate surface proteins

Surface localization can be highly contextual, depending on factors like cell state (Leung et al., 2020; Müller et al., 2021) and cell type (van Oostrum et al., 2019), so a database listing without traceable evidence is of limited use. Our triage agent can scan the whole proteome, but its output is limited to a verdict, a reason code, and a short rationale. This is sufficient to flag a candidate, but not to judge the strength and provenance of the surface localization claim. We therefore created a deep-dive agentic pipeline to assess candidate surface proteins with traceable literature evidence.

The deep-dive cohort consisted of 5,332 surfaceome candidates (Figure 4a): the union of Sonnet and database calls, minus 1,401 proteins found in only one database that the triage agent rated high-confidence non-surface. For each protein, the pipeline returned a structured record backed by per-claim literature citations. It ran in three stages (Figure 4b; Methods): Stage 1, evidence retrieval: surface-specific search seeds drove retrieval across PubMed (Sayers et al., 2022), Europe PMC (Ferguson et al., 2021), and PubTator (Wei et al., 2024). After rule-based removal of reviews, a Haiku agent triaged abstracts to select papers to read, and full text was recovered via a tiered fallback of PMC, Unpaywall (Piwowar et al., 2019, 2018), and DataCite (Brase, 2009). We then used a Haiku trim pass to cut each retrieved paper down to its relevant clips, and a Sonnet pass to select the final evidence into a ledger of verbatim, validated quotes. Trimming and selection were run twice per protein, in two parallel tracks that each produced a ledger. In the Surface Evidence track we kept clips that directly document the protein at the cell surface and the assays used to show accessibility. In the Surface Biology track we kept clips on biological context (expression level and breadth, subcellular localization, and the cell states or lineages in which the protein becomes surface-exposed).

**Figure 4.**
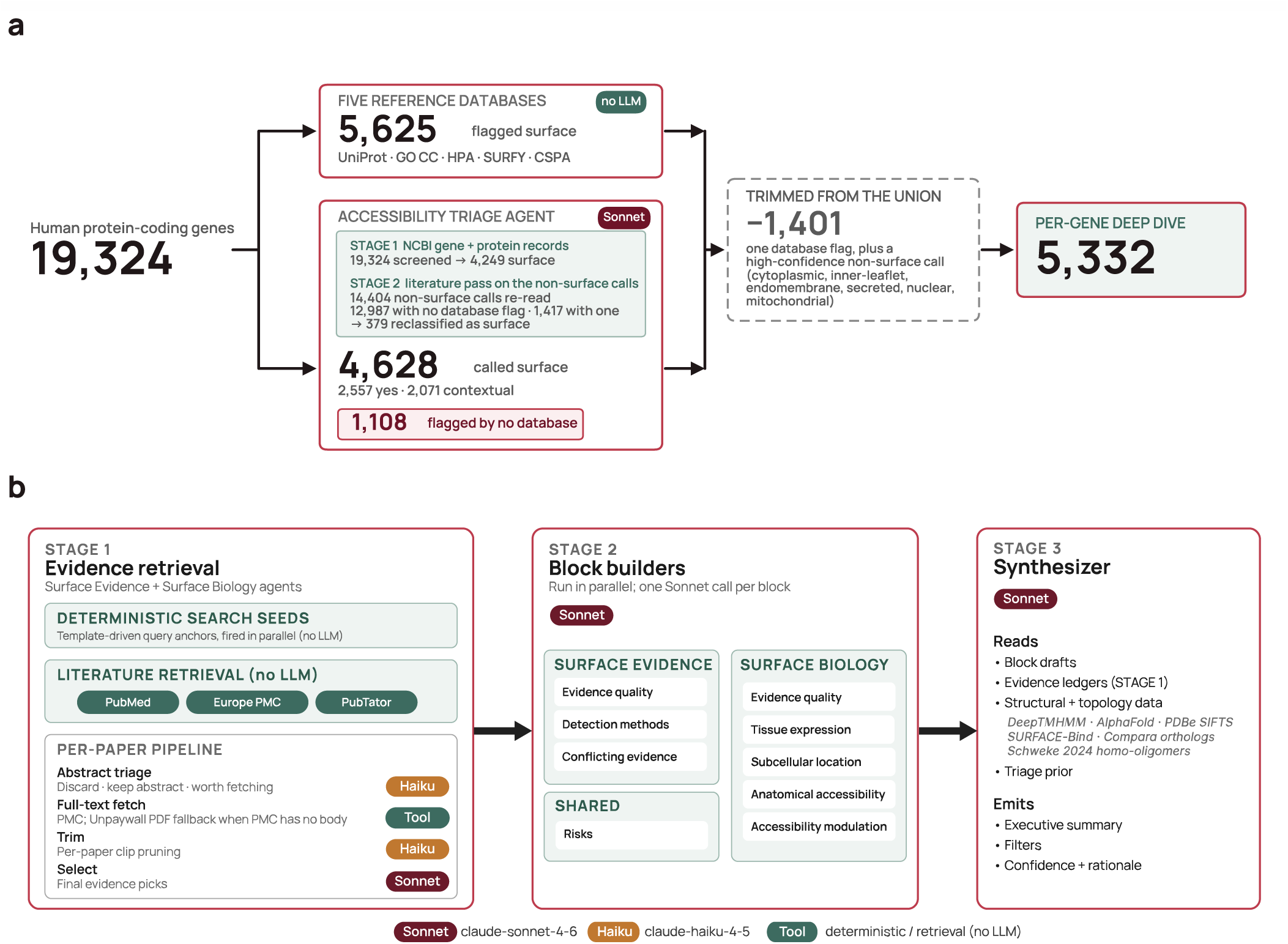
Database and triage calls select 5,332 genes for the deep dive, which runs in three sequential stages, producing one validated record. a) Every protein-coding gene is assessed by the five reference databases and, in parallel, by the two-stage triage agent. Proteins that only one database calls surface, and that the triage agent independently rates high-confidence non-surface with localization to a clear non-surface compartment (noted in the figure), are excluded; the rest of the union (5,332 genes) goes forward to the deep dive. b) The per-gene deep-dive pipeline. Stage 1, evidence retrieval: public databases build a candidate corpus, a per-paper Haiku 4.5 abstract triage selects papers worth fetching, open-access full text is recovered through a tiered fallback, and a Haiku 4.5 trim and Sonnet 4.6 selector yield per-protein evidence sets. Stage 2, block builders (Sonnet 4.6): draft the record’s surface-evidence, surface-biology, and risk sections. Stage 3, synthesis (Sonnet 4.6): the synthesizer reads every section draft plus the Stage 1 evidence set, deterministic features, and the triage verdict, then writes the executive summary, structured filters, and confidence rationale.

Stage 2, block builders: eight Sonnet agents each read only the section of the ledger relevant to their block and answered one defined question (e.g., evidence quality, or combined subcellular location and anatomical accessibility; Figure 4b).

Stage 3, synthesis: one synthesizer call received the block drafts together with the precomputed structural and topology features and the triage verdict. It wrote the executive summary, three overall catalog filters, and an overall confidence call, each with a rationale. The remaining structured filters were filled in deterministically from the block outputs and the precomputed features.

In the viewer, we pair each record’s LLM synthesis and evidence ledger with seven precomputed deterministic features from public structural and orthology resources: membrane topology and ECD length with DeepTMHMM (Hallgren et al., 2022); predicted and experimental structures for the canonical protein, isoforms, close paralogs, and mouse and cynomolgus orthologs from AlphaFold DB (Bertoni et al., 2026) and the PDBe, with orthology from Ensembl Compara (Dyer et al., 2025; Vilella et al., 2009); SURFACE-Bind predicted binding sites (Balbi et al., 2026); and predicted homo-oligomerization (Schweke et al., 2024).

Because our agents can only use papers they can retrieve, we next asked how much of the discovery corpus was actually readable. For 100 randomly selected proteins, discovery returned a median of 235 candidate papers per protein; we sampled 10 papers per protein and attempted full-text recovery. Of these, 89% were recoverable, almost all through PubMed Central (88%); 10% had no open-access copy and under 1% were open access but blocked to automated retrieval. We also tested whether a broader search would help. OpenAlex returned 3.6-fold more papers per protein, but we could read only 49% of them (Figure S10; Supplementary Note 1). Of the rest, 38% were paywalled, 11% were labeled open access but could not be retrieved by automated agents, and 2% were open-access copies that our retrieval missed.

### The deep dive yields a confidence-tiered and configurable surfaceome

We ran the deep-dive pipeline across all 5,332 proteins. The deep dive cost $7,607, or $1.43 per protein; running it on every protein-coding gene would have cost approximately $28,000. Deep-dive accuracy slightly exceeded the triage agent’s, achieving 97.3% accuracy on the 113 SurfaceBench proteins included in the deep dive compared to the triage agent’s 96.8% (Figure S11).

The deep dive did, however, call two benchmark non-surface proteins, ATG9A and RPN1, surface in the low tier, on weak, indirect literature that manual review had missed; we kept the curator’s labels, so both count as errors.

On the three positive-control target sets, the deep dive returned surface calls for 98.3% of ADC antigens, 100% of TCE targets, and 98.4% of viral-entry receptors, closely tracking the triage agent. It differed from triage on only three ADC antigens (COLEC10, FMOD, KLK3), each a secreted or matrix protein rather than a bona fide cell-surface antigen, which the deep dive called non-surface.

The cohort partitioned into 1,783 canonical and 1,252 likely surface calls, and the remaining 2,297 fell in the low, uncertain, and no tiers (Figure 5a). Across the canonical, likely, and low tiers, 1,323 calls carried a cell-type-restricted facet, 434 a cell-state-induced (oncogenic) facet, and 198 a cell-state-induced (other) facet (Figure 5b). Confidence applies to non-surface calls too: of the 1,228 proteins called non-surface, 434 were high-confidence, well-studied proteins with clear evidence of a cytoplasmic, inner-leaflet, endomembrane, or secreted-only location, and 515 were low-confidence, mostly sparsely studied proteins.

**Figure 5.**
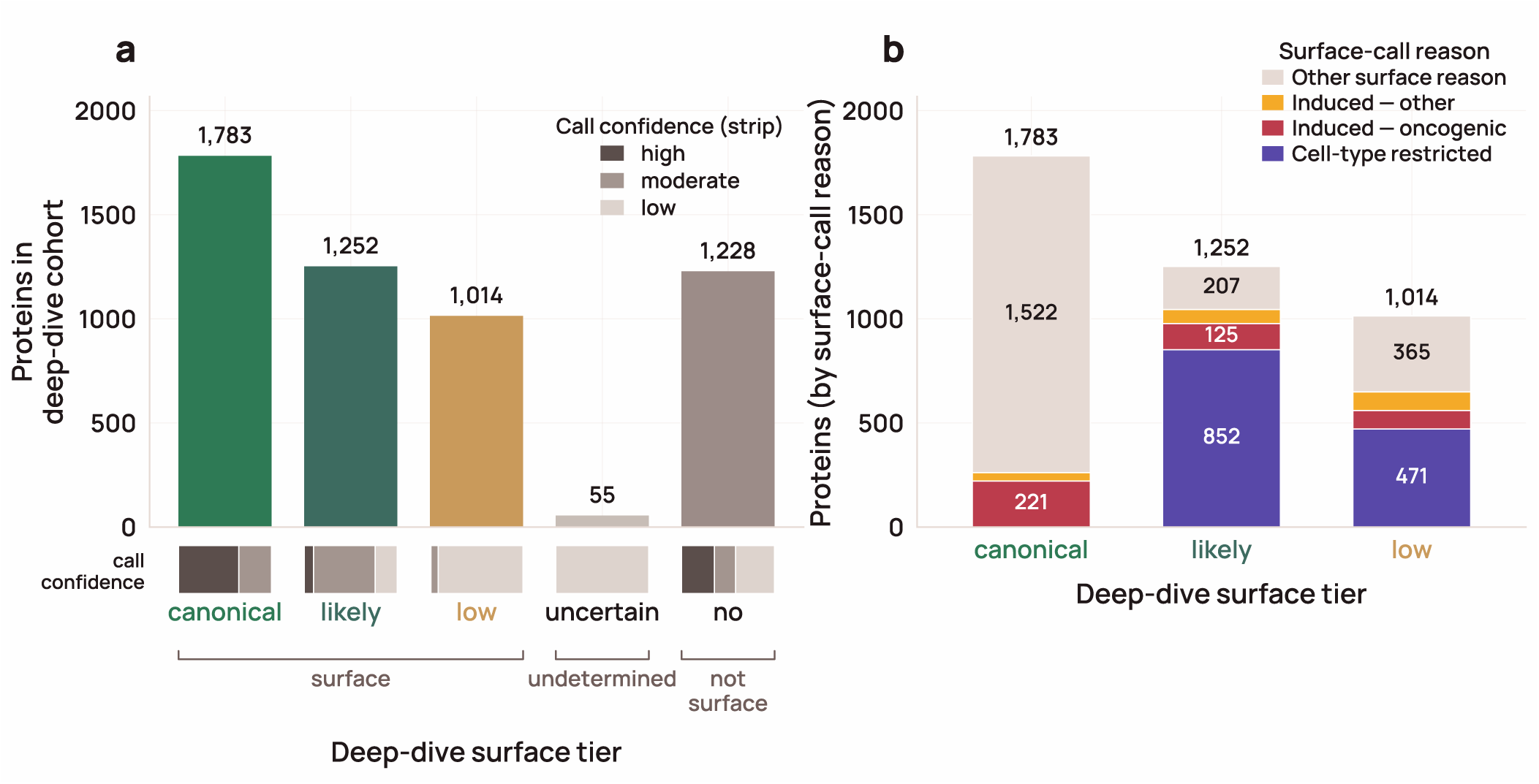
Every deep-dived protein resolves to a surface tier, and the borderline calls carry their own reason for falling short. a) The full deep-dive cohort placed on a five-tier surface scale: canonical (strict surface) and likely (broader), then the below-likely proteins split by the deep dive’s tentative call — low (low or moderate accessibility on weak evidence), uncertain, and no (leaned non-surface). The strip under each bar shows the share of that tier’s calls made at high, moderate, or low confidence. b) Surface-tier proteins from (a) broken down by the deep dive’s surface-call reason: cell-type restricted, cell-state induced (oncogenic or other trigger), or another surface reason (e.g., classical receptor, multi-pass with exposed loops, GPI-anchored, dual localization). Each bar totals the corresponding tier in (a). Canonical proteins were required not to be cell-type restricted but could be highly state-dependent, provided their baseline expression was not reported to be low (see Methods). Counts are over the full deep-dived cohort (n = 5,332). [Figure data and script](https://gist.github.com/beccajcarlson/c2441f8d0314c5524463bc85a3e86612).

The structured tags are most useful for proteins where a single surface label is misleading. One example is SRC, an inner-leaflet kinase that two databases (GO and HPA) list as surface, presumably because of its membrane association. Our record also calls SRC surface, but with high state dependence (disease-induced), direct evidence from only a single method type or source, and a lysosomal-exocytosis mechanism (Delaveris et al., 2026). SRC is therefore recorded as a conditional surface protein and not a constitutive one. CD63, by contrast, is called surface by four of five databases, but the deep dive calls it mostly intracellular: at baseline it is lysosomal and traffics to the surface mainly through activation-induced exocytosis, making it weak as a standalone surface target but potentially credible for bispecific approaches where endolysosomal delivery is desirable (de Goeij et al., 2016).

Across the 5,332 proteins, all with at least one paper found, discovery returned a median of 220 candidate papers per protein (Figure S12). Discovery is deliberately broad: most of these papers mention the protein in passing or study other aspects of its biology. The pipeline kept a median of 10 papers per protein as cited evidence: those flagged in abstract triage as likely to report localization that yielded at least one verbatim supporting quote (Figure 6a–c). We considered an experiment direct evidence if an antibody or probe reached the protein from outside an intact cell, and supporting evidence if it only localized the protein to the plasma membrane without directly demonstrating extracellular accessibility. Of the 3,035 proteins called canonical or likely, 2,203 had at least one direct method, most often a functional surface assay (1,429), flow cytometry (1,186), or surface biotin pulldown (592) (Figure S13).

**Figure 6.**
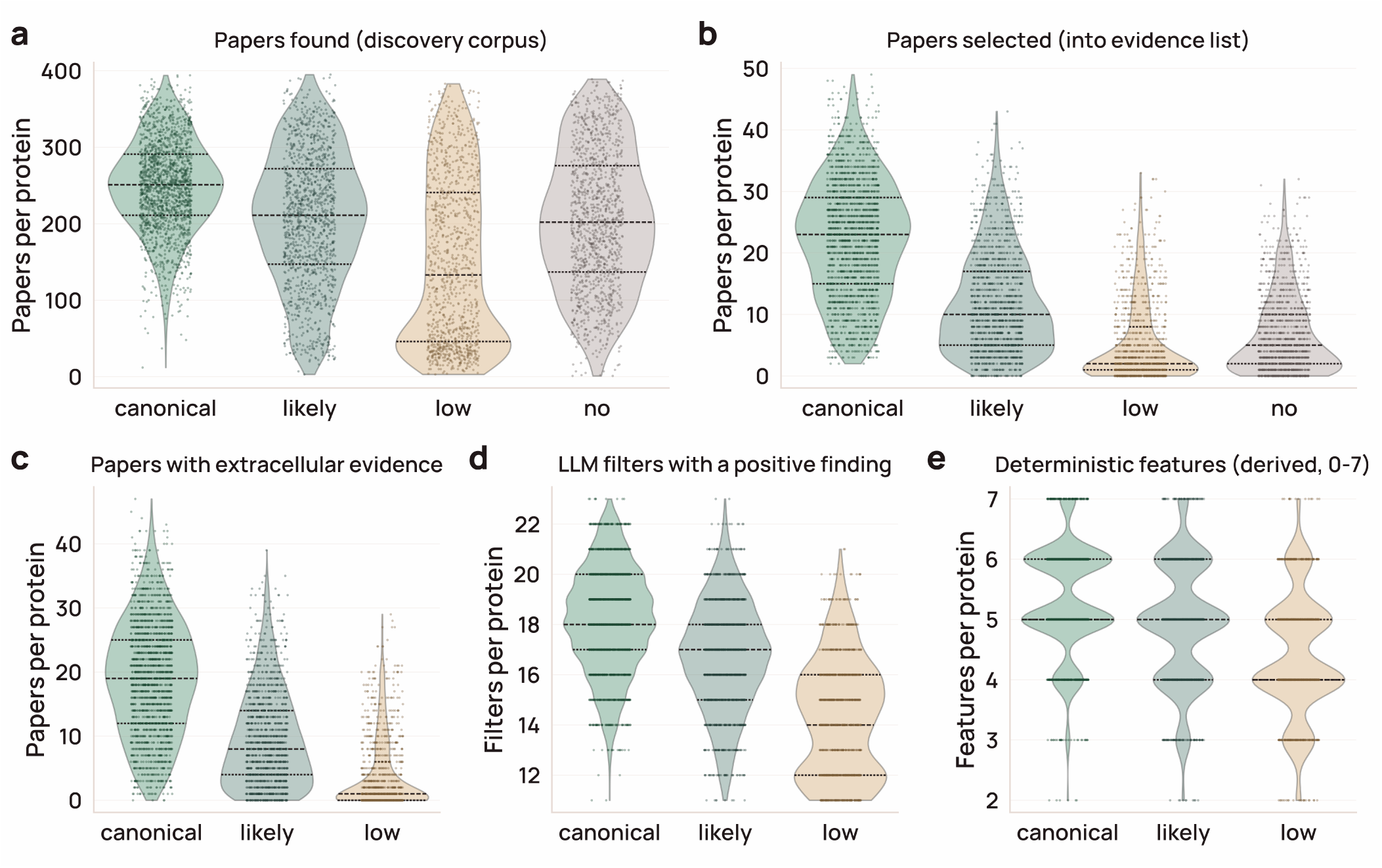
Each deep-dive record is information-dense, and its richness scales with the confidence of the surface call. Five axes of content: a) papers found in discovery, b) papers selected for full-text reading, c) selected papers carrying extracellular evidence, d) structured catalog filters holding a positive, non-default finding (of 24), and e) precomputed deterministic features populated (of 7). Some proteins, for example, lack SURFACE-Bind predictions or known orthologs. Every panel is faceted by deep-dive surface tier — canonical, likely, low, and non-surface (“no”); see Methods. Panels c–e show only the three surface-positive tiers, since these axes summarize the evidence behind a surface call. Each violin is the per-protein distribution for that tier (inner lines: median and 25th/75th percentiles; dots are individual proteins) across the deep-dived cohort, excluding the 55 uncertain-tier proteins, too few to plot (n = 5,277 in panels a and b; 4,049 in panels c–e). [Figure data and script](https://gist.github.com/beccajcarlson/35119ea2bca9585c7245d247334b8c01).

Triage and the deep dive largely agreed on which proteins were surface: 64% of triage-yes proteins were called canonical and 28% likely, 76% of triage-no proteins remained non-surface, and the two stages assigned the same surface-call reason to 50% of proteins (Figure S14).

Each record populates a median of 16 of the 24 catalog filter fields and five of the seven deterministic-feature blocks (Figure 6). Topology and structure are available for all proteins, and orthologs and paralogs for 88% and 87%, respectively, while predicted extracellular SURFACE-Bind binding sites and homomer predictions are positives-only (25% and 19%) (Figure S15).

A low surface tier can reflect how little has been published on a protein rather than its underlying biology. Only 3% of proteins with fewer than 100 discovered papers were canonical, compared with 24% of those with 100–200 papers and 46% of those with 200 or more (Figure S12). Part of this trend is expected, because we required moderate or high evidence density for the canonical tier (Methods). We therefore flagged proteins that had fewer than 100 discovered papers (the 15.8th percentile) but were called surface by UniProt, the best-performing database on SurfaceBench. Of the 538 flagged proteins, the deep dive called 514 surface (21 canonical, 161 likely, and 332 low): a shortlist of probable surface proteins that are under-characterized rather than genuinely low-confidence. Readers can retrieve this list with the “Low-lit + UniProt” filter in the viewer’s catalog table, or through the public API.

Deep-dive records were reproducible across independent runs. We re-annotated 50 randomly drawn proteins from scratch and compared each of the 24 LLM-derived fields with the published record (Figure S16). The five-tier surface scale (no < uncertain < low < likely < canonical) agreed with a linearly weighted κ of 0.85 (95% CI 0.72–0.93). The median κ across the 24 fields was 0.83, which is conventionally considered strong agreement (κ 0.80–0.90) (McHugh, 2012). We then re-ran only the section builders and synthesizer on the original evidence for each record. Tier agreement increased to κ = 0.92 (95% CI 0.84–0.98) and the median field κ to 0.92, so most of the run-to-run variation came from which papers were retrieved and retained, and less from how the model interpreted the same evidence. The four proteins whose surface call changed all had records citing ten or fewer papers, and three of these four calls did not change when we held the evidence fixed. For reference, professional GO curators annotating the same articles chose the identical term in 39–47% of cases (Camon et al., 2005; Van Auken et al., 2014), equivalent to a κ of roughly 0.4.

### An open-source viewer makes the reasoning and evidence behind every surface call inspectable

We built a viewer in which users can filter surface proteins and read the cited evidence behind each call (Figure 7). The viewer, available at surfaceome.deliverome.org, has an accompanying public API and documentation. The viewer code is in the project repository, and the full records served by the API are deposited on Zenodo with the repository code (Carlson and Smallegan, 2026a, 2026b). Because records will continue to be updated, we give each record numbered revisions and periodically freeze the full catalog into a numbered data release on Zenodo (see Methods), so that a call can be cited and retrieved later as it was at the time.

**Figure 7.**
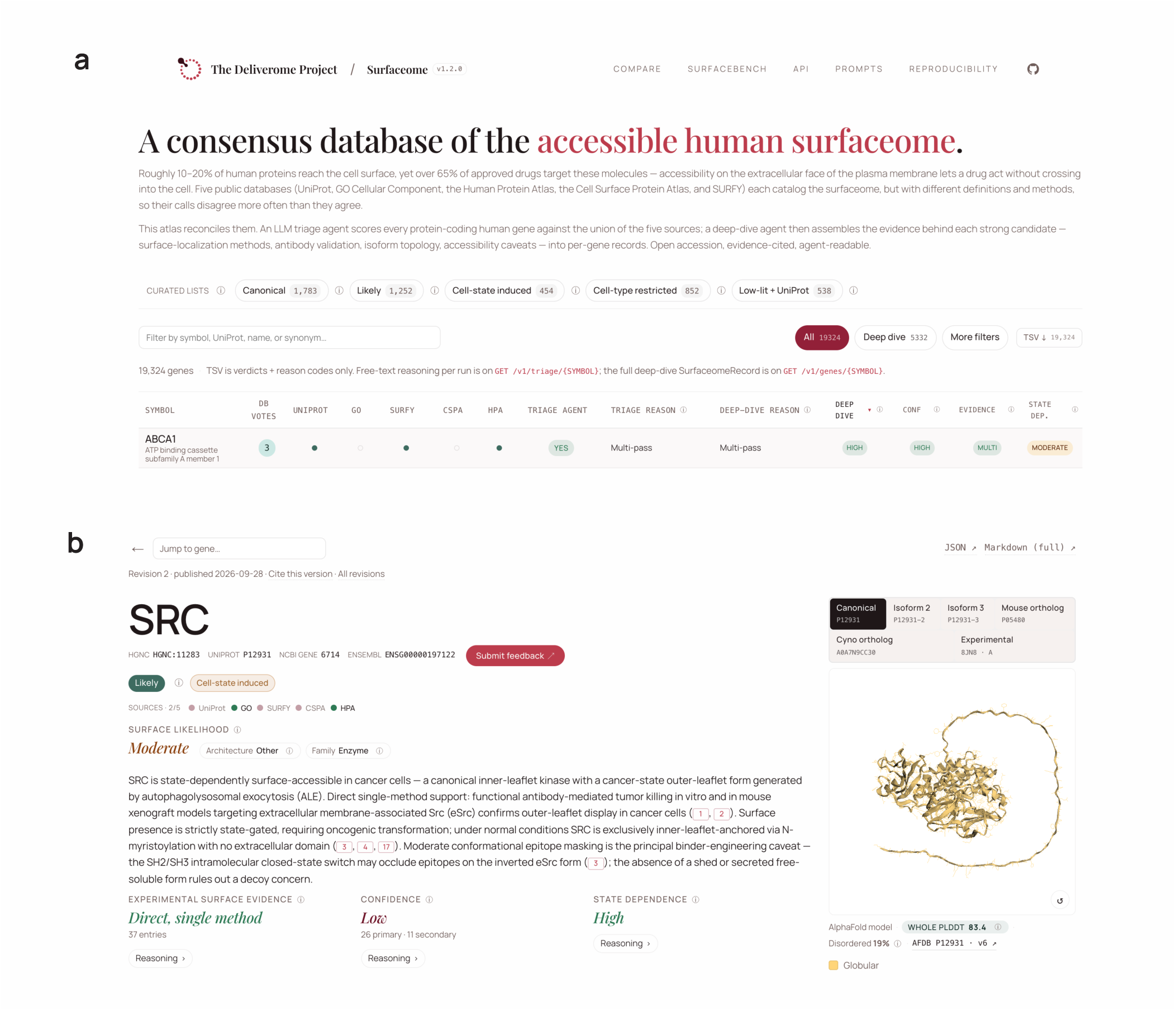
A fully open-source surfaceome viewer. a) The viewer homepage, showing a table of all proteins with reference database membership, triage LLM calls, and deep-dive LLM calls. b) For proteins with a deep-dive record, a detailed page is available with both LLM-derived and deterministic results.

On the landing page, we show all 19,324 protein-coding genes in a sortable table with the five source database calls and the triage verdict, and for deep-dived genes the deep-dive verdict, confidence, evidence grade, and state dependence. Any record field can be used as a filter. We included presets for the canonical and likely tiers and for the cell-state-induced and cell-type-restricted groups in Figure 5, but the individual fields (e.g., evidence grade or ECD accessibility class) can also be filtered separately if a user wants to examine different thresholds. We did not make the low-literature flag a tier, so it can be added to any query (for example, to find canonical or likely proteins that also have little published literature).

For proteins with a deep-dive record, the record page has the surface-localization verdict and confidence, the graded attributes with their supporting verbatim excerpts, and the deterministic features. It also lists the current revision, what changed since the last revision, the data release, and a permanent link.

In the Compare view, a user can enter a list of candidate proteins and see the surface call, tier, and evidence for each one, similar to enrichment tools like Enrichr (Chen et al., 2013; Xie et al., 2021). The viewer also has pages for the agent prompts, the SurfaceBench benchmark with calls from each model, and the reproducibility materials. Finally, each record has a feedback form where users can flag errors or missing information, with an option to make the report public. We welcome broader suggestions or contributions at.

## DISCUSSION

Existing surface protein databases agree on only 188 proteins (Figure 1), and our results suggest that much of this disagreement reflects context rather than errors. Each database derives its calls from a different kind of evidence and records them as fixed labels, but the conditions under which a protein reaches the surface are mostly described in the free text of individual papers. An LLM triage agent with no external tools classified SurfaceBench proteins more accurately than any of the five databases or their ensemble (Figures 2, S5). UniProt, with an optimized cutoff, handled clear cases similarly well, while the LLM’s advantage lies mainly in detection of context-dependent proteins. Proteome-wide, 1,021 of the 1,108 triage candidates absent from all five databases were contextually surface (Figure 3).

Grounding these calls in the full-text literature through a follow-up deep-dive agentic pipeline preserved accuracy while adding details behind each overall verdict. The result is a configurable surfaceome rather than a single list, with cell-type and cell-state facets that let users assemble the set of surface proteins that best fits their question. Because all calls also carry confidence and metadata on evidence found, the records separate proteins with clear evidence of a non-surface location from those that are simply understudied, a distinction lost when a protein is merely absent from a database.

Since records will change as literature, models, and sources are updated, they are versioned so analyses can pin to a numbered release; the results reported here correspond to release 1.3.0. New literature accrues quickly: in each of 2024 and 2025, roughly 60% of deep-dived proteins gained at least one newly linked publication in NCBI’s gene2pubmed. Poorly studied proteins gained new papers less often (29–40% per year), but all four surface calls that changed on re-annotation were in records citing ten or fewer papers (Figure S16), so we expect to refresh our database up to once annually, prioritizing understudied proteins.

The cost of building resources like this one is also falling quickly. When we re-ran the triage benchmark on the later Claude 5.5 models, the smallest, Haiku 5.5, came within a point of our production Sonnet 4.6 configuration at a small fraction of the cost (Figure S3). Newer models also brought new failure modes. Opus 5.5’s safety classifiers declined to answer about 5% of calls, all on ordinary surface proteins. Pipelines built on LLMs should therefore treat a refusal as an explicit outcome rather than a silent gap, and re-benchmark each model upgrade rather than assume it improves accuracy.

The viewer intentionally says little about whether a protein has been targeted therapeutically or has suitable binding sites. Therapeutic modalities, from antibodies and antibody–drug conjugates to CAR-T cells and radioligands, each place different demands on a target, such as epitope size and location, surface copy number, or internalization. While we include SURFACE-Bind sites as a starting point, targetability is a modality-specific question beyond the scope of this viewer. We likewise leave clinical information to resources such as Open Targets (Buniello et al., 2025). More broadly, we built the catalog to be combined with other data; through the public API, users can join it with their own internal data or with other public resources such as OpenAntigens, which suggests antigen constructs for human cell-surface and secreted proteins (Teixeira et al., 2026).

A caveat specific to a resource built by LLM agents is that the pipeline scales expert judgment but does not yet replace it. Accurate records required sustained expert oversight, with extensive testing and manual review to catch failure modes without overfitting to the cases we had flagged. Every call therefore carries a graded confidence, and low, uncertain, and no calls, especially those resting on sparse literature, should be read as provisional. The per-claim evidence and public feedback mechanism are meant to make errors easy to spot and correct.

Coverage of any such database is bounded by what machines can read. In an audit of a random sample of surface proteins, 89% of papers from our targeted discovery were readable in full, almost all via PubMed Central; a broader OpenAlex search found 3.6-fold more papers, but only 49% of them were readable (Figure S10; Supplementary Note 1). The bot-accessible corpus thus collapses largely to PubMed Central, so our database reflects not the state of the science but the portion of it that machines are currently permitted to read. Coverage is also limited by what has been measured at all. Systematic screens are beginning to make these measurements directly, quantifying plasma membrane expression across 766 GPCRs (Tedman et al., 2026) and across thousands of variants of individual channels and receptors (Coyote-Maestas et al., 2022; Howard et al., 2025; Mighell and Lehner, 2025). Such screens typically test overexpressed constructs in one cell line, while the literature captures native cell types and states; integrating both into records like ours would let each inform the other.

Surface accessibility is necessary but not sufficient for delivery. To carry a therapeutic into a target cell, a surface protein must also be present at sufficient copy number and internalize the therapeutic once bound; we call this subset of the surfaceome the deliverome. Neither property is well captured in the literature: internalization data are far sparser than surface-expression data, and surface abundance is hard to infer from existing measurements. Protein abundance correlates poorly both with RNA abundance (Gygi et al., 1999; Liu et al., 2016; Wang et al., 2019; Wu et al., 2026) and with compartment-specific protein abundance such as surface abundance (George et al., 2026b), and quantitative global proteomics data scaled across subjects are only recently becoming available (Ding et al., 2025; Fang et al., 2025; Knol et al., 2025; Malmström et al., 2025; Yue et al., 2026). These datasets still under-sample surface proteins. Closing these gaps will require physical experiments as well as better computational methods. We envision LLM agents coupled directly to these experiments in the future, identifying where evidence is thin or contradictory and prioritizing what to measure, with the resulting data feeding the next round of predictions. At Deliverome Bio, we are generating internalization and surface-abundance data toward this goal and intend to share them promptly via this viewer and public repositories.

The approach we used here is not specific to surface proteins. A cheap proteome-wide triage followed by evidence-anchored deep dives suits any question whose observations are scattered across papers but only partly captured in databases, such as context-dependent localization to intracellular compartments, regulators of protein trafficking or activity, or ligand–receptor pairings. Although the databases we compared against did not outperform LLMs, without them we could not have built a benchmark or measured performance at proteome scale. As LLMs grow more capable, curated databases will matter more as the ground truth that agents are measured against. We offer this resource as an addition to that infrastructure, with every call, quote, and prompt released in machine-readable form so that the next scientist, and the next agent, can build on it.

## DATA AVAILABILITY

Data are permanently available via Zenodo (https://zenodo.org/records/20805383) as well as live via our API. All figure gists containing input data and self-contained code for plotting are stably deposited with Software Heritage. The SWHID list can be found here: https://github.com/Deliverome-Project/accessible-surfaceome/blob/v1.3.0/data/analysis/figures/figure_swhids.json.

## CODE AVAILABILITY

All code, including the web viewer code, is available at https://github.com/Deliverome-Project/accessible-surfaceome under an MIT license. Code is also permanently available via Zenodo (https://doi.org/10.5281/zenodo.22116981).

## METHODS

### Candidate universe construction

Because each source defines surface membership differently, we evaluated alternative inclusion rules for each database against SurfaceBench and kept the most accurate variant for each, so that every database was compared at its best (Figure S1). Two of five rules changed (UniProt and CSPA); the other three were retained as published.

### UniProt

For the UniProt database, we initially categorized a protein as surface if its subcellular_locations field contained any of four strict terms (“Cell surface”, “Apical cell membrane”, “Basolateral cell membrane”, “GPI-anchor”) or if it had at least one extracellular topology feature (“feature_topo_extracellular” > 0), giving 3,175 proteins. “Cell membrane” alone was excluded because it includes cytoplasmic-side and lateral-membrane proteins. After recalibration (Figure S1), we also accepted proteins with a transmembrane domain or signal peptide (“feature_transmembrane” > 0 or “feature_signal_count” > 0), giving 3,894 proteins (SurfaceBench accuracy from 78% to 90%) and recovering 719 GPI-anchored and single-pass receptors that UniProt curates with “Cell membrane” only rather than the stricter surface terms.

### SURFY

For SURFY, we accepted the published binary classifier output (“surfy_is_surface” = 1), yielding an initial set of 2,886 proteins. Tighter classifier thresholds (>0.5, >0.7, or >0.9) reduced overall accuracy without offsetting gains in specificity on SurfaceBench, so we retained the authors’ binary call. SURFY is not independent of UniProt and CSPA: its positive training set comprised 910 α-helical transmembrane proteins found in at least two of CSPA’s high-confidence set, UniProtKB/Swiss-Prot entries with the “cell membrane” keyword, and five-star COMPARTMENTS plasma-membrane proteins, and its published surfaceome adds 130 GPI-anchored proteins taken from UniProt annotations.

### CSPA

For CSPA, we initially flagged proteins categorized as high confidence or putative (excluding “unspecific”), giving 1,286 proteins. After recalibration, we kept high-confidence only, which gave comparable recall on SurfaceBench with substantially better specificity (accuracy from 48% to 52%) and an optimized set of 1,039 proteins.

### GO CC

For GO CC, we initially flagged a protein as surface when it carried at least one non-electronic annotation for a surface CC term (allowing experimental, curated, or sequence-based evidence). Proteins with only electronic-inference (IEA) annotations were excluded, giving an initial set of 2,023 proteins. Admitting IEA-only annotations raised benchmark accuracy from 42.2% to 48.3% but added 534 proteins, so we kept the initial rule (Figure S1).

### HPA

For HPA, we flagged a protein as surface when it was categorized as “Plasma membrane” or “Cell Junctions” and the antibody was in the Enhanced, Supported, or Approved reliability tier (excluding Uncertain), giving 2,107 proteins. No alternative cutoff tested improved benchmark accuracy.

### Treatment of source absence

The union of all five databases contained 5,680 proteins (5,546 genes, the unit counted in Figure 1) under the initial cutoffs and 5,993 under the optimized cutoffs: 391 added by UniProt’s expanded rule and 78 dropped by CSPA’s tighter rule. These counts are of UniProt accessions. The deep-dive cohort is counted per gene over the 19,324 triaged protein-coding genes, where the optimized union contained 5,625 genes; adding the 1,108 genes called surface by triage alone and removing the 1,401 trimmed genes gives the 5,332-gene cohort. Absence from UniProt, SURFY, GO CC, or CSPA counted as a “not surface” vote. SURFY is a bounded predictor rather than a proteome-wide annotation: its features are computed per topological domain, so it is defined only over the 7,903 proteins with a determinable membrane topology, with labels published for the α-helical subset. Because the trained classifier is not distributed, its coverage cannot be extended, so we scored an absent SURFY label as non-surface, matching what a user of the published resource obtains. HPA’s coverage is limited by antibody availability and validation (∼13,600 proteins total), so absence from HPA counted as an abstention and was excluded from accuracy (98 of the 147 SurfaceBench proteins were present in HPA).

### Benchmark curation

To evaluate the triage agent and reference databases on the hardest-to-categorize part of the surfaceome, we curated a 147-protein benchmark (SurfaceBench) enriched for proteins where the five databases disagreed. A single curator reviewed the literature for each protein and assigned one of three ground-truth verdicts (surface, contextually surface, or non-surface), along with a reason drawn from a 19-category taxonomy (Table S1). SurfaceBench was hand-selected rather than randomly sampled. Most proteins (138 of 147) received surface calls from one to four of the five databases, and we chose them to span positive (approved-drug targets, GPI-anchored proteins, GPCRs), borderline (state-induced, dual-localized, or partner-attached), and negative classes (secreted proteins, intracellular drug targets, and proteins on the wrong membrane face or in the wrong compartment). The remaining nine received the same call from all five databases. We deliberately included a few such proteins, both easy controls that every database calls surface and hard cases that every database calls incorrectly, and recalibrating the database cutoffs after the benchmark was assembled (Figure S1) moved a few more to unanimous agreement. Per-protein verdicts, reasons, rationales, and database votes are available on the [SurfaceBench page](https://surfaceome.deliverome.org/benchmark/).

### Benchmark evaluation and statistics

For scoring, “yes” and “contextual” verdicts were collapsed into a single surface class. Each classifier (the triage agent and each database) was scored correct when it assigned a ground-truth surface protein to the surface class and a non-surface protein to “no”. Each model and prompt combination was run in three replicates per protein. Per-replicate verdicts (available through the viewer and API) were aggregated to a majority verdict per model and prompt combination for per-protein comparisons; reported accuracies are the mean of per-replicate accuracies, with SEM across replicates. Paired comparisons between classifiers used an exact two-sided McNemar test on per-protein correctness, with the majority verdict across replicates. Because few proteins were discordant in any comparison, these tests have low power, and a non-significant result does not establish equivalence. We also scored database ensembles calling a protein surface when at least *k* of the five databases did, for *k* = 1 (union) through 5 (unanimous). Each was scored under the same soft-credit rule (Figure S5). No ensembling condition exceeded the accuracy of the Sonnet agent alone.

### Positive-control target sets

As independent positive controls, we assembled three lists of validated surface proteins: antibody–drug conjugate (ADC) targets, T-cell engager (TCE) targets, and viral-entry receptors.

### ADC targets

The ADC list (n = 234) is the union of targets from TheraSAbDab, Open Targets ADC annotations, and ADCdb. For TheraSAbDab (Raybould et al., 2020), we retrieved entries designated as antibody–drug conjugates and removed non-cytotoxic conjugates mislabeled as ADCs. We then added targets of drugs annotated as ADCs in Open Targets release 26.06 (Buniello et al., 2025). Because these drugs have mechanism-of-action records for both the antibody and its cytotoxic payload, we kept only records whose target was the antibody’s cell-surface antigen. For ADCdb (Shen et al., 2023), the largest and least thoroughly validated database, we applied a text filter to the antigen name and functional description to remove intracellular, nuclear, and secreted protein families that had been incorrectly included. Multi-target records were mapped to their primary antigen only. After we joined all three sources, Open Targets contributed no antigens beyond those already in the other two. A few borderline secreted or extracellular-matrix proteins from ADCdb still survived filtering (for example, fibromodulin, fibronectin, and cathepsin D). Notably, ADCdb targets have heterogeneous evidence levels, and many are preclinical, still-speculative targets rather than validated surface proteins.

### T-cell engager targets

The TCE list (n = 30) was also derived from TheraSAbDab (Raybould et al., 2020). Because TheraSAbDab has no TCE annotation, we took the targets of antibodies whose format was bispecific, trispecific, or tetraspecific and whose target list included CD3E. The final set comprises 28 disease-target antigens plus the CD3D and CD3E chains of the engaged T-cell CD3 complex.

### Viral-entry receptors

The viral-entry-receptor list (n = 62) comprises human cell-surface viral-entry receptors from Expasy ViralZone (Hulo et al., 2011). Protein receptors were resolved to UniProt accessions and then HGNC identifiers; non-protein entries (e.g., glycan and glycolipid receptors) and non-human accessions were excluded.

The final lists are in the source data for Figure S7: https://gist.github.com/beccajcarlson/3ab5d-f749b576912959c75fe7013d78c and stably deposited at https://archive.softwareheritage.org/swh:1:rev:30631354d617077d5c2ea942a43ee345c385fc2a;path=/positive_control_db_coverage_bars.tsv.

### Triage agent

The triage agent classified each protein as yes, contextual, or no for plasma-membrane surface accessibility, with a single reason drawn from a fixed 19-category set and a brief rationale. Eight Claude models (Haiku 4.5 and 5.5; Sonnet 4.6, 5, and 5.5; Opus 4.8, 5, and 5.5) were run across up to four context conditions per model for all 147 SurfaceBench proteins: (i) gene symbol only; (ii) a resolved-identifier condition pairing the NCBI summary with a canonical identifier table (HGNC-approved name, UniProt accession, aliases, previous symbols, HGNC gene groups, and CD designation); (iii) condition ii plus an on-demand web-search tool; and (iv) literature anchoring, in which PubMed evidence for cell-surface accessibility was retrieved in advance and appended to the prompt. All variants share verdict definitions, reason taxonomy, pre-“no” checklist, and output schema, differing only in context; the prompts are archived with the code release (prompts at v1.3.0).

### Proteome-wide two-stage triage

All 19,324 protein-coding genes in the cohort were triaged in two stages. In stage one, a tool-free triage agent (Sonnet 4.6 with NCBI gene context) classified each protein in a single replicate. Stage two was a single PubMed-augmented pass (condition iv above) that re-read stage-one “no” calls, so that a documented surface form could overturn the stage-one call. The pass covered two groups of genes: database-negative proteins, absent from every reference database (n = 12,987), and single-database proteins, the 1,417 high-confidence “no” calls carried by exactly one database, which were re-checked before the conservative trim was applied (see Cohort-scale execution, Candidate universe). A protein reclassified as yes or contextual was kept; in all, stage two reclassified 379 proteins as surface (Figure 4a). The two-stage sweep recovered 1,108 candidate surface proteins that were absent from all five reference databases.

### Deep-dive annotation pipeline

#### Literature retrieval

All literature sources are free to query; we used an identifying user-agent and did not bypass robots.txt.

#### Discovery

Each protein’s candidate search list was the union of three queries: a) NCBI PubTator3 named-entity recognition (NER)-anchored search, in the form @GENE_<symbol> <topic terms>, returning papers in which NCBI’s NER tagged the protein as a subject entity, not as an incidental mention (Wei et al., 2024); b) NCBI gene2pubmed cross-references, surfaced via Europe PMC’s EXT_ID disjunct, returning National Library of Medicine–curated paper-to-gene links (Sayers et al., 2022); and c) Europe PMC keyword search, which provided topic-anchored backfill against paired gene-synonym and surface-method clauses (Ferguson et al., 2021). On a random 100-protein test cohort, this union returned a median of 234.5 papers per protein, ranging from ∼12 for orphan proteins to ∼412 for well-studied receptors.

#### Candidate filtering

Papers without an abstract were not considered further. When a batch of newly discovered candidates numbered at least 50, rule-based filters removed reviews by publication type (except those in high-quality review journals) and, by title, drug-development reviews and atlas or proteome-wide screens, and the remaining batch was capped at 160 papers, keeping the most recent. On three heavily studied proteins these rules retained 94–100% of papers that contributed evidence. A Haiku 4.5 agent then read each abstract and either discarded the paper (it studies a different protein or mentions this one only in passing), kept the abstract alone (it already states the surface-biology finding), or marked it for full-text reading; if the full text could not be recovered, the abstract was used instead. A Haiku trim pass cut each retrieved paper to its relevant passages, explicitly retaining evidence against surface localization (for example, primarily intracellular localization or no detectable surface signal), and a Sonnet pass selected the final evidence, tagging each claim as supporting, refuting, or ambiguous. Papers removed before the trim pass were not audited for contradicting evidence.

#### Full-text recovery

For each candidate paper selected for evidence extraction, full-text retrieval was attempted through three tiers, moving to the next on a miss. (1) PMC JATS XML, queried first through NCBI E-utilities and then falling back to Europe PMC’s mirror; PMID-only papers were linked to PMC via NCBI eLink’s pubmed_pmc linkname. JATS was preferred because it preserves section structure, which otherwise must be reconstructed from PDF layout. (2) Unpaywall, which resolves the DOI to a list of open-access (OA) locations; up to four PDF URLs were attempted, ranked by quality (publisher publishedVersion above repository acceptedVersion). PDFs were streamed under an 80 MB cap and parsed by pdfplumber with gutter-based two-column detection and font-aware heading recognition. (3) DataCite landing-page scrape, where the DataCite REST API returns the landing URL and the OA license; the landing page’s <meta name=“citation_pdf_url”> tag provides the article PDF URL. The same 80 MB cap applied. Each clip records its recovery tier and, when available, its OA license.

#### OpenAlex

We evaluated OpenAlex (Priem et al., 2022) as an alternative discovery source on the same 100-protein cohort under matched 21-axis search parity and did not adopt it, because it added few reachable on-topic papers at higher LLM cost and its free-tier rate limit was incompatible with a proteome-wide sweep (Supplementary Note 1; Figure S10).

#### Deep-dive pipeline architecture

Each protein in the candidate universe (in at least one reference database or called surface by the two-stage triage, minus the 1,401-protein trim; see Candidate universe) was processed by a deep-dive pipeline that returns a single structured record (SurfaceomeRecord) with per-claim literature citations and an immutable set of precomputed deterministic features. The pipeline calls the Anthropic Messages API directly. End-to-end cost was a median of $1.32 per protein (mean $1.43; 10th–90th percentile $0.50–$2.47) at the Sonnet 4.6 tier, with a median wall-clock time of ∼9 minutes (550 s; 10th–90th percentile 3.8–17 min).

The pipeline runs in three stages. First, a plan_trim_ select step issues two paired Sonnet calls, one for each track (Surface Evidence and Surface Biology), that read the triage record, the saved literature corpus, and the protein’s identifier bundle. These decide which evidence subsets each downstream block builder receives, bounding the cost of the fan-out. Second, eight block builders populate the record’s nine sections in two waves: six run concurrently (methods, contradictions, expression, subloc_anatomical_combined, accessibility_modulation, and biological_context_grade, where the combined builder fills both the subcellular-localization and anatomical sections), and two run afterward because they need inputs the first wave does not have. The evidence_grade builder uses the methods builder’s per-claim annotations to assign the evidence grade: direct_single_method, direct_ multi_method, supportive_but_indirect, conflicting, or weak; and the risks builder reads the combined Surface Evidence and Surface Biology evidence together with the deterministic features, which are fetched after the first wave, to identify surface-accessibility risks. Each methods row records the method family and subclass, permeabilization, expression system (endogenous or overexpressed), and whether the experiment shows direct surface accessibility, supports plasma-membrane localization, supports membrane association only, or reports expression only. For Figure S13, rows were grouped into eight method categories by subclass where family and subclass disagreed (for example, cell-surface capture filed under a biotinylation or glycoproteomics family), and only direct and supporting rows were counted. Third, a synthesizer reads all nine sections and the deterministic features and writes the executive_summary block, including the headline surface-accessibility verdict, the surface_call_reason enum value drawn from TriageReason, and overall confidence.

Every record is validated against the SurfaceomeRecord Pydantic schema. A draft that fails validation re-runs from the last completed stage; the plan_trim_select output is checkpointed, so a retry costs only the failed sub-call (∼$0.05) rather than the full plan_trim_ select phase. The schema separates LLM-graded fields from deterministic features: the deterministic_ features region is read-only to every LLM call, and a Pydantic cross-validator on the synthesizer’s draft rejects any agent-written value in this region. Before synthesis, three deterministic passes are applied to the risks builder’s output: the ECD-accessibility class is set from the DeepTMHMM ECD length, a homo-oligomerization prediction is attached from the Schweke atlas, and a secreted form is flagged when an isoform has no transmembrane helix but keeps at least 30 extracellular residues. After synthesis, deterministic consistency rules drop headline risks that the structured risk fields do not support, withdraw a secreted-form risk that has no citation and no corroborating localization or fractionation evidence, and reconcile state dependence, surface accessibility, and the surface-call reason with one another. Validated records are published as JSON snapshots and to the public Cloudflare D1 database that backs the API.

#### Deep-dive scoring

Calls are made per gene: a protein counts as surface if any documented proteoform reaches the extracellular face, such as a membrane-bound splice form, the transmembrane precursor of a shed ectodomain, or a secreted protein held at the surface by a partner. Forms that could compete with or mask the surface form (shed or secreted forms, isoform decoys) are recorded as risks, conflicting isoform-level evidence is recorded as a contradiction, and membrane topology is reported for every UniProt isoform. Each protein with a deep-dive record was assigned to exactly one of five mutually exclusive surface tiers (canonical, likely, low, uncertain, or no), spanning a spectrum from gold-standard surface evidence to a negative call (proteins lacking a deep-dive record were excluded). These tiers are separate from each record’s confidence field, which rates certainty in the call in either direction, and from the evidence grade, which rates only the strength of evidence for surface localization. Tiers are deterministic predicates over the record’s structured scores; the two positive tiers match the catalog’s presets.

The evidence grade used for tiering is the synthesizer’s summary grade (evidence_grade_summary), which takes the same values as evidence_grade. A protein qualified as likely if its grade was direct_multi_ method, direct_single_method, or supportive_but_indirect, or was weak with a structurally unambiguous surface_call_reason (classical_surface_receptor, multipass_with_exposed_loops, gpi_anchored, or tissue_restricted_surface); surface_specificity ∈ {surface_dominant, mixed, mostly_intracellular}; and surface_accessibility ∈ {high, moderate}. It qualified as canonical (a stricter subset of likely) if its surface_ call_reason was neither tissue_restricted_surface nor lysosomal_exocytosis, its grade was direct_multi_ method, direct_single_method, or supportive_but_indirect, and it satisfied surface_specificity ∈ {surface_ dominant, mixed}, surface_accessibility ∈ {high, moderate}, confidence ∈ {high, moderate}, evidence_density ∈ {high, moderate}, and either state_dependence ∈ {low, moderate, unclear} or expression_level ∈ {moderate, high}. Canonical was evaluated first, so the plotted likely tier excludes canonical proteins.

Proteins failing the likely predicate (mostly those with weak or conflicting experimental evidence) were split into three tiers by their tentative surface_accessibility call: low (surface_accessibility ∈ {low, moderate}; a positive-leaning “maybe surface” call below the likely bar), uncertain (surface_accessibility = uncertain; no directional signal), and no (surface_accessibility = no; a negative-leaning call). These tiers reflect the direction of the lean, not evidence strength; weak-evidence proteins populated all three, most falling in low. Because they rest on thin literature, these tiers are provisional.

#### Cohort-scale execution

##### Candidate universe

Proteins were carried forward to deep-dive annotation if they were in ≥1 of the five reference databases or called positive by the two-stage triage (union rule). We then applied a conservative trim, removing proteins called high-confidence non-surface by triage and supported by only one database (1,401 proteins). Every trimmed protein had first been re-checked with PubMed in stage two and was still called “no”: 1,364 of the 1,417 in the single-database group (the other 53 were reclassified and kept), plus 37 supported only by UniProt’s expanded rule, which had been database-negative under the initial cutoffs and so were re-checked in the database-negative group.

##### Deep-dive scaling

Each protein in the candidate universe was annotated independently, parallelized across on-demand cloud workers (Modal; up to ∼200 concurrent).

##### Run-to-run reproducibility

To measure reproducibility, we drew 50 proteins uniformly at random from the 5,332 deep-dive records, in two disjoint batches of 25, and re-analyzed each in two ways (Figure S16). In a full re-run, the pipeline was re-executed from scratch with the same prompts and models, three months after the published run, so the comparison also captures new literature and changed search results. In a fixed-evidence replay, each record’s published evidence was reused and only the section builders and synthesizer were re-run. For each of the 24 LLM-derived filter fields, the binary surface call, and the five-tier surface scale, we computed Cohen’s κ between the re-analysis and the published record, using linear weights for ordered fields and unweighted κ for nominal fields and yes/no flags (Sim and Wright, 2005). State dependence and co-receptor dependency were scored on their ordered levels only, excluding records called unclear or unknown. Confidence intervals were 95% percentile intervals from 5,000 paired bootstrap resamples of proteins, and agreement was interpreted using previously published bands (McHugh, 2012). Re-analyzed records are not part of the published catalog; all three versions of each sampled record (published, full re-run, and replay) are available as labeled reproducibility replicates through the public API (/v1/replicates) and in the Zenodo data record.

### Deterministic features

Each record carries seven precomputed deterministic features from public structural and orthology resources.

#### 1. Cross-species orthology

Mouse and cynomolgus one-to-one high-confidence orthologs come from Ensembl Compara via BioMart REST (release 115) (Dyer et al., 2025; Vilella et al., 2009). One-to-many and many-to-many relationships are dropped at ingest; the Ensembl orthology-confidence flag is read directly from BioMart. Per retained ortholog we record the Ensembl gene ID, symbol, and full-length and ECD-restricted %-identity to human canonical.

#### 2. Paralogs

Paralogs come from Ensembl Compara’s within-species homology table (same BioMart source, release 115). Per protein, we retain the top-N paralogs ranked by ECD %-identity to canonical; DeepTMHMM topology and sequence are attached only to close paralogs whose full-length identity to canonical is ≥80% (Edfors et al., 2018). More distant paralogs carry identity statistics only. The 80% floor follows Edfors et al., who found that above this identity a single-target antibody is generally unachievable.

#### 3. Isoforms

We enumerate UniProt-annotated alternative isoforms from the topology sweep. Because Compara does not pair splice variants of one gene, we compute pairwise identity directly using the same BLOSUM62 helper used for paralogs and orthologs, retaining full-length and ECD-restricted %-identity plus an ECD BLOSUM62-positive similarity. Each isoform’s per-residue topology is projected onto the canonical residue axis so all variants align in the viewer.

#### 4. Membrane topology

We run DeepTMHMM 1.0.24 (Hallgren et al., 2022) across four cohorts — human canonical, annotated alternative isoforms, mouse orthologs, and cynomolgus orthologs — with every distinct paralog UniProt accession added to the human canonical cohort so that ECD %-identity is computable for every human–paralog pair regardless of full-length divergence. DeepTMHMM is preferred to UniProt’s curated topology, which is sparser for less-studied proteins and lags isoform-specific updates. From the per-residue calls we compute the extracellular-domain length in residues (ecd_length_ residues) and, before synthesis, bin it into one of five accessibility classes: none, minimal (<30 residues), small (30–59), moderate (60–199), or large (≥200). The synthesizer prompt lists the same thresholds, but the rendered value is always the deterministic one.

#### 5. AlphaFold structure

For each canonical UniProt accession we record its AlphaFold DB (Bertoni et al., 2026) entry — entry ID and version, the mean pLDDT over the ECD, the fraction of ECD residues below pLDDT 50 as a disorder proxy, and the model’s CIF, PDB, and PAE file URLs. We also record one representative experimental structure per canonical UniProt accession, chosen from the PDBe SIFTS UniProt→PDB best_structures mapping by highest coverage, then best (lowest) resolution (Dana et al., 2019).

#### 6. Candidate binding sites

For proteins covered by the SURFACE-Bind atlas (Balbi et al., 2026), we ingest the published per-site MaSIF-scored binding-patch arrays — site identifier, anchor residue, buried surface area in Å², an Eisenberg-style patch hydrophobicity score (positive = lipid-facing; negative = solvent-exposed), and per-site α-helical and β-strand seed counts (the number of de novo binder backbones passing MaSIF scoring) — and the per-protein summary counts that the synthesizer reads. SURFACE-Bind covers 1,649 of the 2,886 predicted surface proteins Balbi et al. scored; proteins absent from the table are marked has_data=false.

#### 7. Homo-oligomerization

Homo-oligomer state comes from the Schweke 2024 AF2-derived homomer atlas (Schweke et al., 2024), which predicts whether each human protein assembles into a homo-multimer in its native context.

### Topology classification and benchmark-composition statistics

Each protein was assigned to exactly one of eight mutually exclusive topology classes derived deterministically from its UniProt keyword annotations (transmembrane-helix count, signal-peptide count, lipidation count, and the GPI-anchor, intramembrane, G-protein-coupled-receptor, prenylation, myristoylation, and secreted keywords) together with its DeepTMHMM per-residue topology prediction. Assignment followed a fixed priority order — GPI-anchored, then intramembrane, then 7TM GPCR (the dedicated GPCR keyword with at least five predicted helices), then generic multi-pass TM (≥2 helices), single-pass TM (exactly one helix), signal-only/likely-secreted (a signal peptide with no TM and no anchor), inner-leaflet lipidated (prenyl or myristoyl anchor with no TM and no signal peptide), and finally no-TM/no-signal — so that, for example, a GPI anchor took precedence over a stray annotated helix. The nine categories reported in Figures S8 and S9 were seven of these classes (all except intramembrane), namely GPI-anchored, 7TM GPCR, multi-pass TM (non-GPCR), single-pass TM, likely secreted, inner-leaflet lipidated, and no-TM/no-signal, plus two overlapping features: glycosylation site (presence of any UniProt-annotated glycosylation feature) and TM without signal peptide (the DeepTMHMM TM-only class). Proteins lacking UniProt keyword or DeepTMHMM data were treated as missing rather than negative.

### Web viewer and public API

#### Viewer

The catalog viewer is a standalone Next.js 16 application (React 19, Node 24), statically exported and hosted on Cloudflare Pages at surfaceome.deliverome.org. Each protein page has a reader-feedback form; moderated public entries are shown on the protein page via /v1/feedback/public.

#### Public API

Programmatic access is provided by a Cloudflare Worker at api.deliverome.org/surfaceome/, with read-only access to the public D1 database (apart from reader-feedback submissions) and open CORS. Resource endpoints cover the annotated-protein list and per-protein SurfaceomeRecord (/v1/genes, /v1/genes/{symbol}), the whole-cohort catalog with per-protein database flags and canonical triage verdict (/v1/catalog), per-gene and per-model-and-prompt triage detail (/ v1/triage/{symbol}, /v1/triage/{symbol}/{model}/{variant}), the SurfaceBench truth and per-model call matrix (/v1/benchmark, /v1/benchmark/matrix), and ortholog lookups (/v1/orthologs/{symbol}). Long-format TSV exports of the triage and benchmark tables (/v1/ triage/export.tsv, /v1/benchmark/export.tsv) support bulk reanalysis and are archived on Zenodo (Carlson and Smallegan, 2026a). Live API documentation is available at https://surfaceome.deliverome.org/api/.

Record history and data releases. Starting after release 1.0.0, the Worker’s served response is archived each time a record is published. The archive covers the record JSON, the per-protein evidence ledger, and the Markdown export, each stored content-addressed by SHA-256. A new revision is written only when one of the three content hashes changes. Hashes exclude serve-time timestamps, so unchanged content does not create revisions. Revisions are numbered from 1 per protein, and each records its publication time, schema version, and prompt-corpus version, plus which components changed. Archived bodies are immutable and served at /v1/genes/{symbol}/revisions/{n} (with /evidence and .md variants). A numbered data release freezes the current revision of every protein. Releases are listed at /v1/releases, record their GitHub tag and Zenodo version DOI, and serve each protein as it stood in that release at /v1/releases/{version}/genes/{symbol}. Each release takes the repository’s semantic version and is deposited on Zenodo, with its records, as a new version of the data record.

## ACKNOWLEDGMENTS

We thank Chance Bowman, Willow Coyote-Maestas, Nirmit Damania, Seymour de Picciotto, Frank Gao, Dann Huh, Kang Yong Loh, Aashish Manglik, Andreas Metousis, Rob Oslund, Evangelia Petsalaki, Max Salm, Austin Tedman, Josh Tycko, Priya Veeraraghavan, and Charles Warren for thoughtful review and feedback on this manuscript.

AI was used to assist in writing the Methods section and editing the rest of the manuscript text. The authors reviewed and take responsibility for the final content.

## AUTHOR CONTRIBUTIONS

Rebecca J. Carlson: Conceptualization, Methodology, Investigation, Software, Validation, Formal analysis, Data curation, Visualization, Project administration, Funding acquisition, Supervision, Writing – original draft, Writing – review & editing.

Michael J. Smallegan: Methodology, Investigation, Software.

L. Robert Hollingsworth: Project administration, Funding acquisition, Writing – review & editing.

## Supplementary information

**Supplementary Table 1.** 19-category reason taxonomy for surface protein triage agent verdicts. Reason codes assigned alongside each verdict (5 surface, 5 contextually surface, 8 non-surface, plus “other”).

| SURFACE | CONTEXTUALLY SURFACE | NON-SURFACE |
| --- | --- | --- |
| **classical_surface_receptor**<br>Single-pass transmembrane protein | **tissue_restricted_surface**<br>Surface form exists only in specific tissues / cell types / context | **cytoplasmic**<br>Soluble cytoplasmic; no membrane association |
| **multipass_with_exposed_loops**<br>Multi-pass TM | **dual_localization**<br>Stable PM pool with dominant non-PM | **nuclear**<br>Nuclear-resident |
| **gpi_anchored**<br>GPI-anchored on the outer leaflet | **stable_surface_attachment**<br>Secreted protein stably attached post-translationally | **endomembrane_resident**<br>Intracellular compartment only; no documented PM access |
| **stable_complex_partner**<br>Protein stably on the surface as part of an anchored complex (e.g., B2M) | **cell_state_induced**<br>Reaches the outer leaflet only in a non-baseline cell state | **mitochondrial_internal**<br>Mitochondrial matrix or matrix-facing inner membrane |
| **extracellular_face_protein**<br>Any other architecture with an explicit extracellular face | **lysosomal_exocytosis**<br>Lysosomal TM protein that reaches the PM during lysosomal exocytosis | **secreted_only**<br>Secreted with no stable surface anchoring |
|  |  | **inner_leaflet_anchored**<br>Lipidated on the cytoplasmic (inner) face of the PM |
|  |  | **pmhc_only_intracellular**<br>Protein body intracellular; documented surface is just MHC-presented peptides |
|  |  | **nuclear_envelope**<br>Inner / outer nuclear membrane only |

**Supplementary Figure 1.**
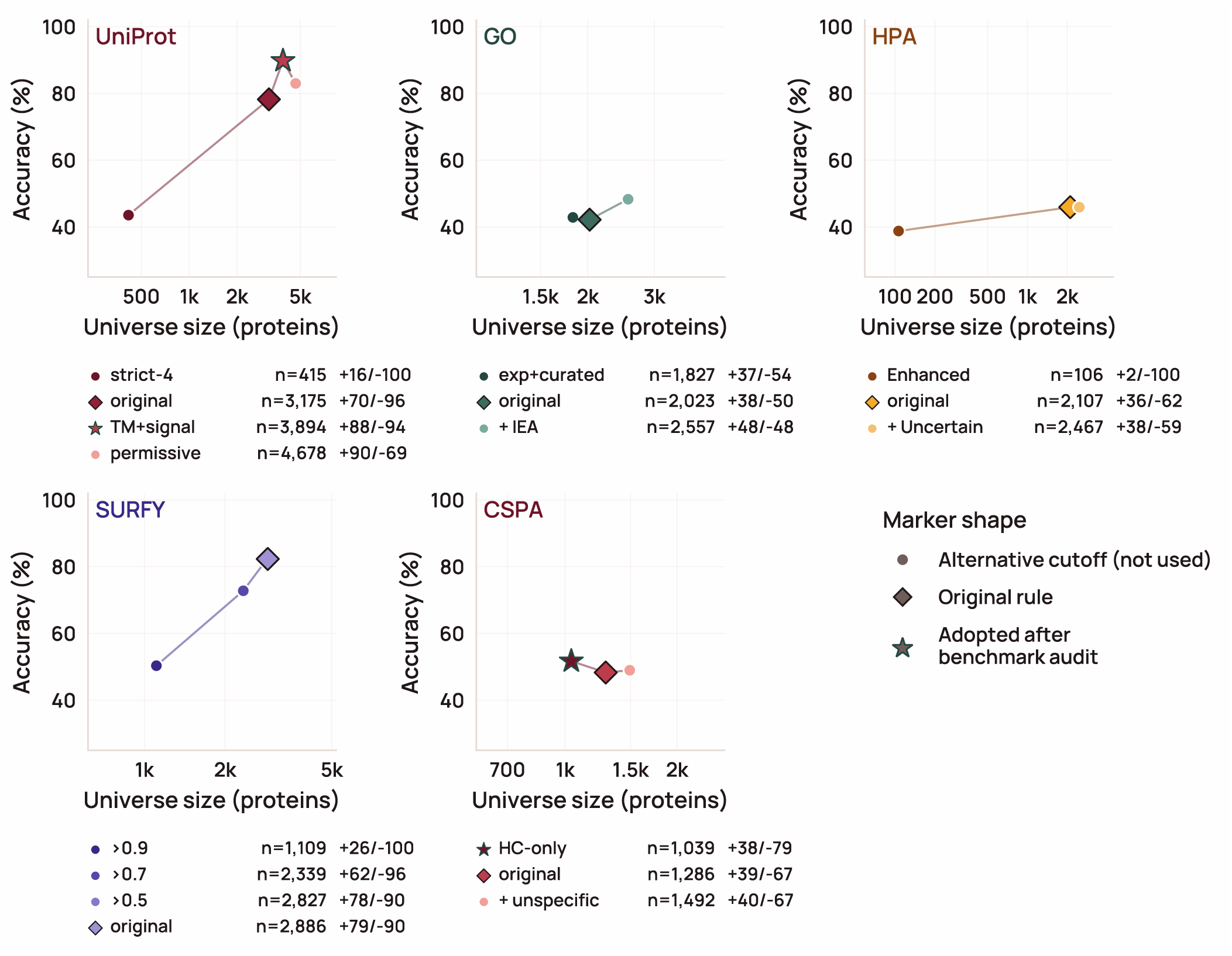
Re-deriving each source database’s surface-inclusion rule against SurfaceBench changes which proteins two of the five sources admit, while the other three are left unchanged. Each panel shows one source database; points are alternative inclusion rules of increasing permissiveness, plotted by the number of proteins each would admit (horizontal axis, log scale; further left is stricter) against accuracy on SurfaceBench (vertical axis). Points are annotated with sensitivity and specificity. The original rule and the rule adopted after the benchmark audit are marked. UniProt was loosened by additionally accepting transmembrane- and signal-peptide-annotated entries, recovering GPI-anchored and single-pass receptors it had been missing with little loss of specificity. CSPA was tightened by restricting it to high-confidence detections only, improving specificity at comparable recall. Human Protein Atlas and SURFY were already at their optimum and left unchanged; GO was left unchanged because its accuracy gain did not justify the extra proteins admitted. [Figure data and script](https://gist.github.com/beccajcarlson/f9319af882e372194bd30640c0cbf2ed).

**Supplementary Figure 2.**
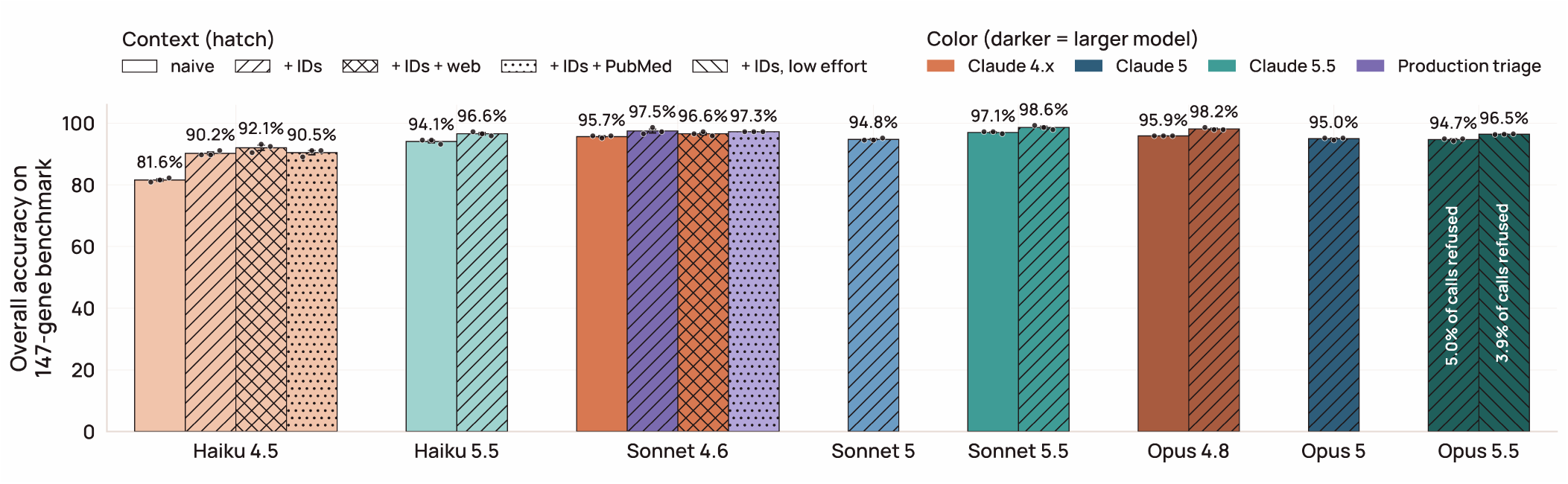
Supplying protein-coding gene identifiers, but not web search or pre-fetched PubMed abstracts, drives within-tier accuracy gains in language-model triage on the 147-protein benchmark. Bars show mean soft-credit verdict accuracy across replicates. Eight Claude models (Haiku 4.5 and 5.5; Sonnet 4.6, 5, and 5.5; Opus 4.8, 5, and 5.5) are compared under up to four conditions: gene symbol alone (naive); symbol plus a cached gene summary and stable HGNC/UniProt identifiers (+IDs); identifier context plus an on-demand web-search tool; and identifier context plus PubMed abstract excerpts retrieved in advance and appended to the prompt. Opus 5.5 was also run with identifiers at low reasoning effort. Three replicates per protein per condition (points; error bars SEM). Refused calls (Opus 5.5 only) are excluded from accuracy, and the share refused is printed on the bar. Color indicates model generation. The two lavender bars are the configurations the production pipeline runs: Sonnet 4.6 with identifiers for the genome-wide triage sweep, and the same model with PubMed abstracts for the stage-two PubMed pass. [Figure data and script](https://gist.github.com/beccajcarlson/9c765ed9673d7bd845c3ac091ad2204d).

**Supplementary Figure 3.**
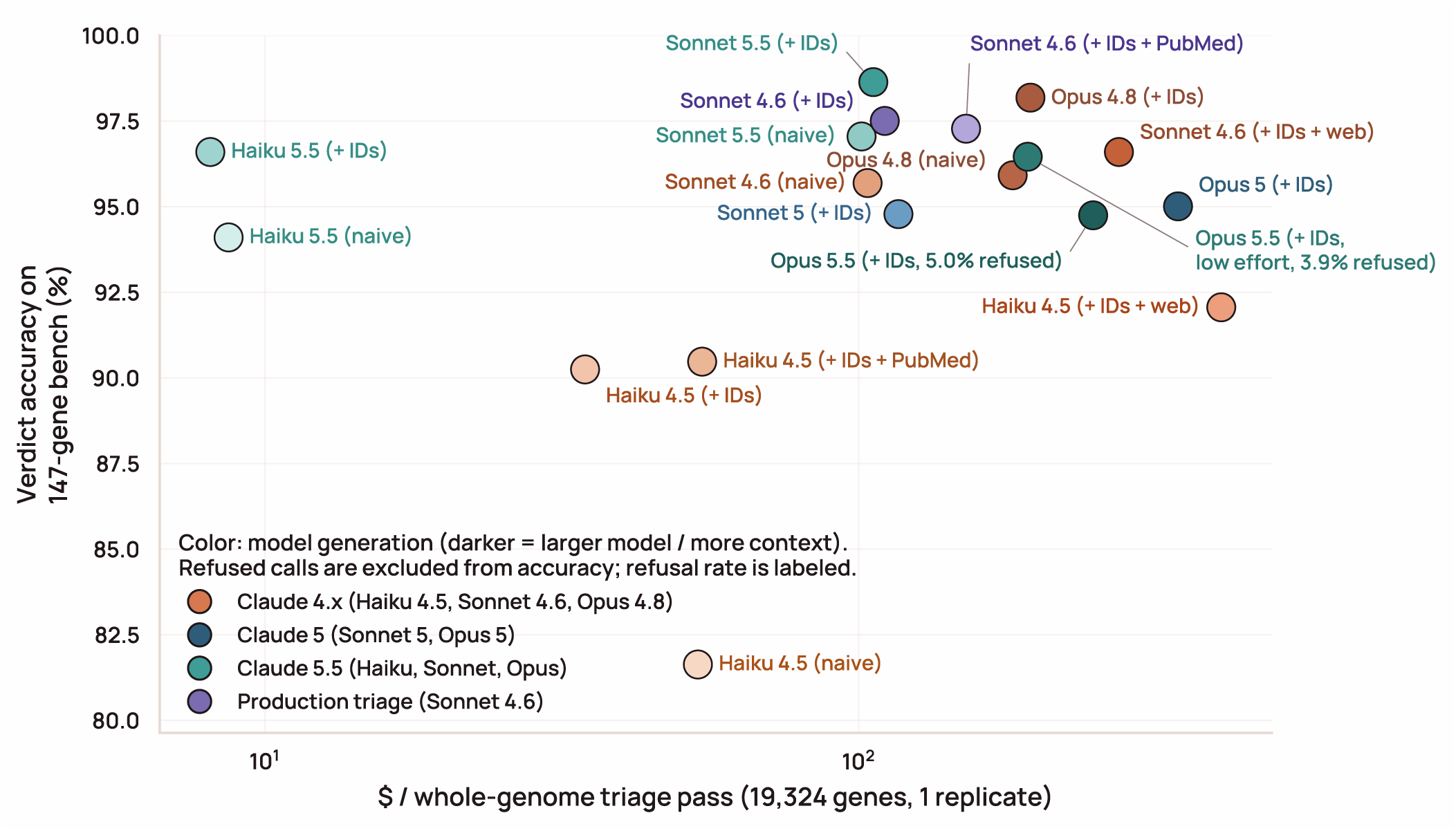
Across model tiers and amounts of supplied context, triage accuracy saturates near 98% well before cost does, so the cheapest configuration reaching that plateau is the economical production choice. Each point is one model–context combination, as in Supplementary Figure 2. The horizontal axis is the projected cost of triaging all 19,324 protein-coding genes once (log scale); the vertical axis is mean soft-credit accuracy on SurfaceBench. Cost is built from average tokens sent and returned per call at Anthropic’s October 2026 list prices, with shared instructions charged once at the first-use rate and reused cheaply, and each web search billed at one cent. Moving from Sonnet 4.6 to Opus 4.8 costs roughly 1.8 times as much for 0.7 percentage points, and adding a search tool costs more than moving to Opus — the two web-search configurations are among the three most expensive points on the chart. Production uses Sonnet 4.6 with identifiers, which came within 0.7 percentage points of Opus 4.8 at the lowest cost among the models available when the pipeline was built; of the later Claude 5.5 models, Haiku 5.5 came within a point of it at about a fourteenth of the cost, and Sonnet 5.5 was the most accurate configuration tested. Refused calls are excluded from accuracy and the refusal rate is labeled (about 5% for Opus 5.5 at its default reasoning effort and 4% at low effort; no other model refused). Color indicates model generation. The two lavender points are the production triage configuration and the PubMed-abstract variant used for the stage-two PubMed pass. [Figure data and script](https://gist.github.com/beccajcarlson/d7f764d2de288ae31cf44173bc396d41).

**Supplementary Figure 4.**
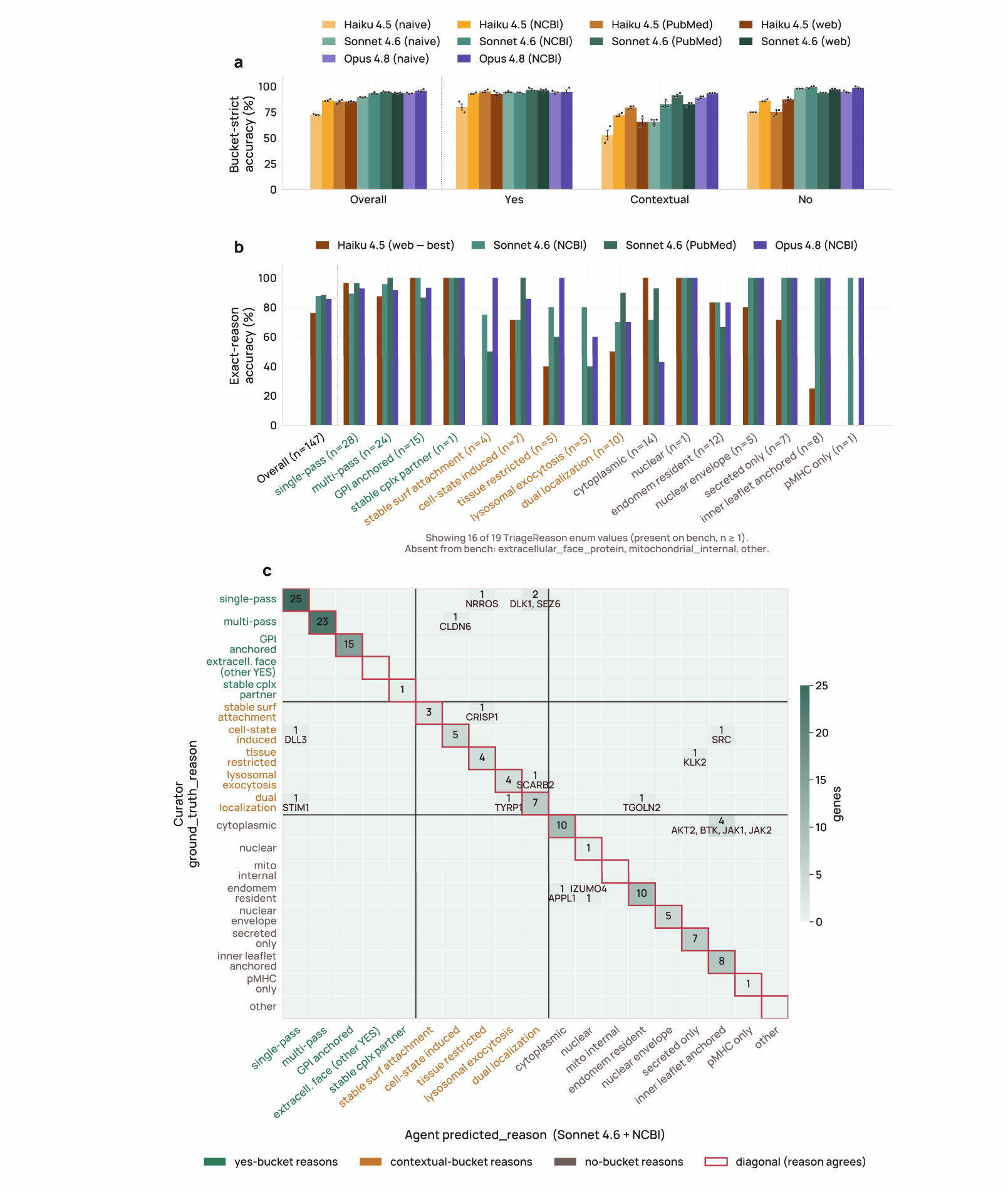
The triage agent’s localization reasoning matches the manual curator on 87.8% of SurfaceBench proteins, with only 3 of 18 residual disagreements moving a protein between surface and non-surface. Curator and agent reasons are drawn from the same closed set (Supplementary Table 1), grouped into yes, contextual, and no buckets. a) Bucket-strict accuracy across all model-and-configuration cells (Haiku 4.5, Sonnet 4.6, Opus 4.8), with per-replicate dots and SEM; a Yes-versus-Contextual mismatch gets no credit. b) Per-reason exact agreement for four frontier configurations (best Haiku, Sonnet with identifiers, Sonnet with PubMed, best Opus), ordered Yes to Contextual to No. c) The Sonnet 4.6 plus NCBI confusion matrix, blocked by bucket; the largest off-diagonal is a within-No-bucket swap that leaves the surface verdict unchanged. [Figure data and script](https://gist.github.com/beccajcarlson/c98fe6646ba8b967767d072657da31b7).

**Supplementary Figure 5.**
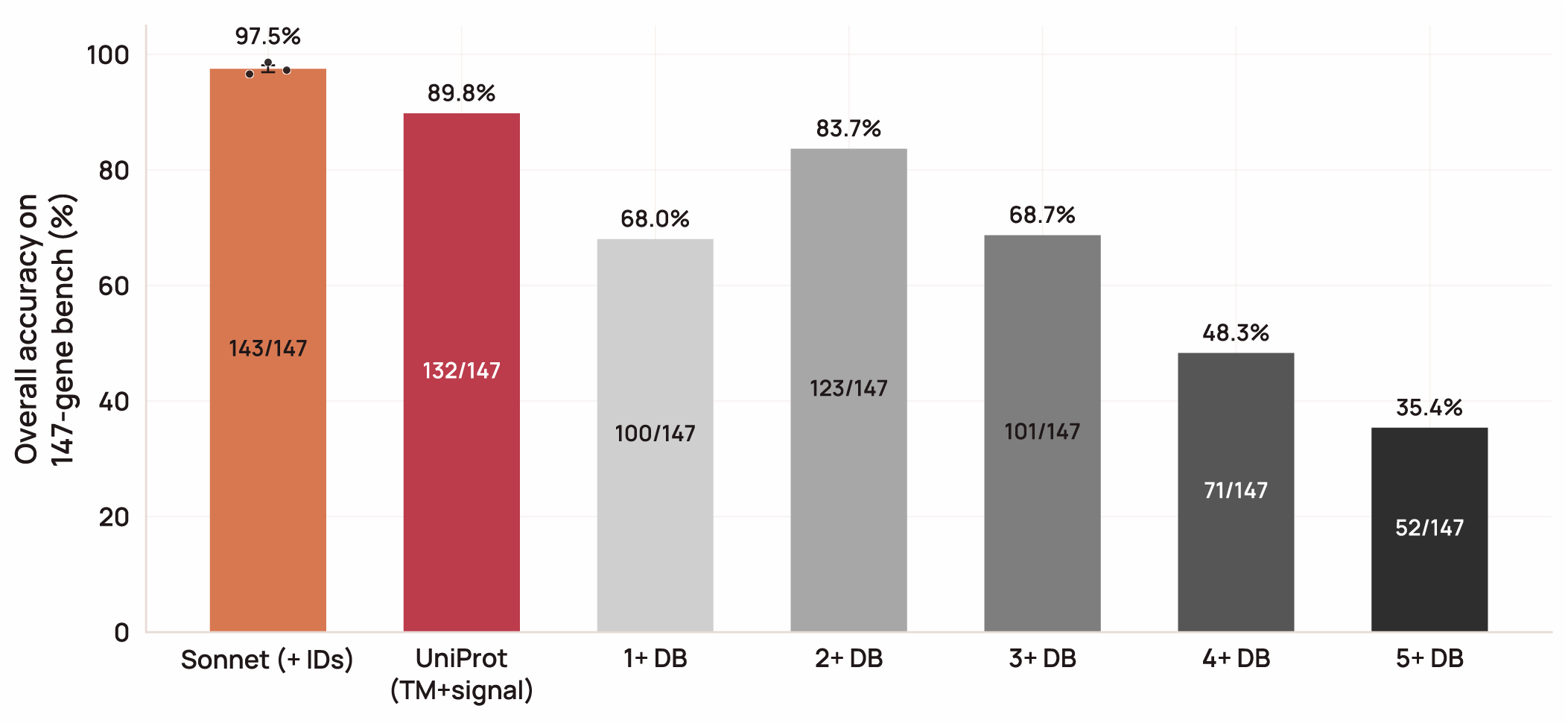
Ensembling the five gold-standard surface databases does not close the accuracy gap to the Sonnet 4.6 triage agent. Soft-credit accuracy on SurfaceBench is shown for seven predictors: the production triage agent (Sonnet 4.6 with identifiers; mean of three replicates, dots; error bar SEM), the single best-performing database (UniProt under its transmembrane- and signal-peptide-expanded rule), and five ensemble predictors that call a protein surface when at least one (the union), two, three, four, or all five databases do. Each database (UniProt, GO cellular component, HPA, SURFY, CSPA) is applied under its SurfaceBench-calibrated inclusion rule (Supplementary Figure 1). Accuracy peaks when two databases must agree (83.7%) and falls on either side. No ensemble matches the best single database (89.8%) or the triage agent (97.5%). [Figure data and script](https://gist.github.com/beccajcarlson/0104308c239fe49d91d82a1007632b27).

**Supplementary Figure 6.**
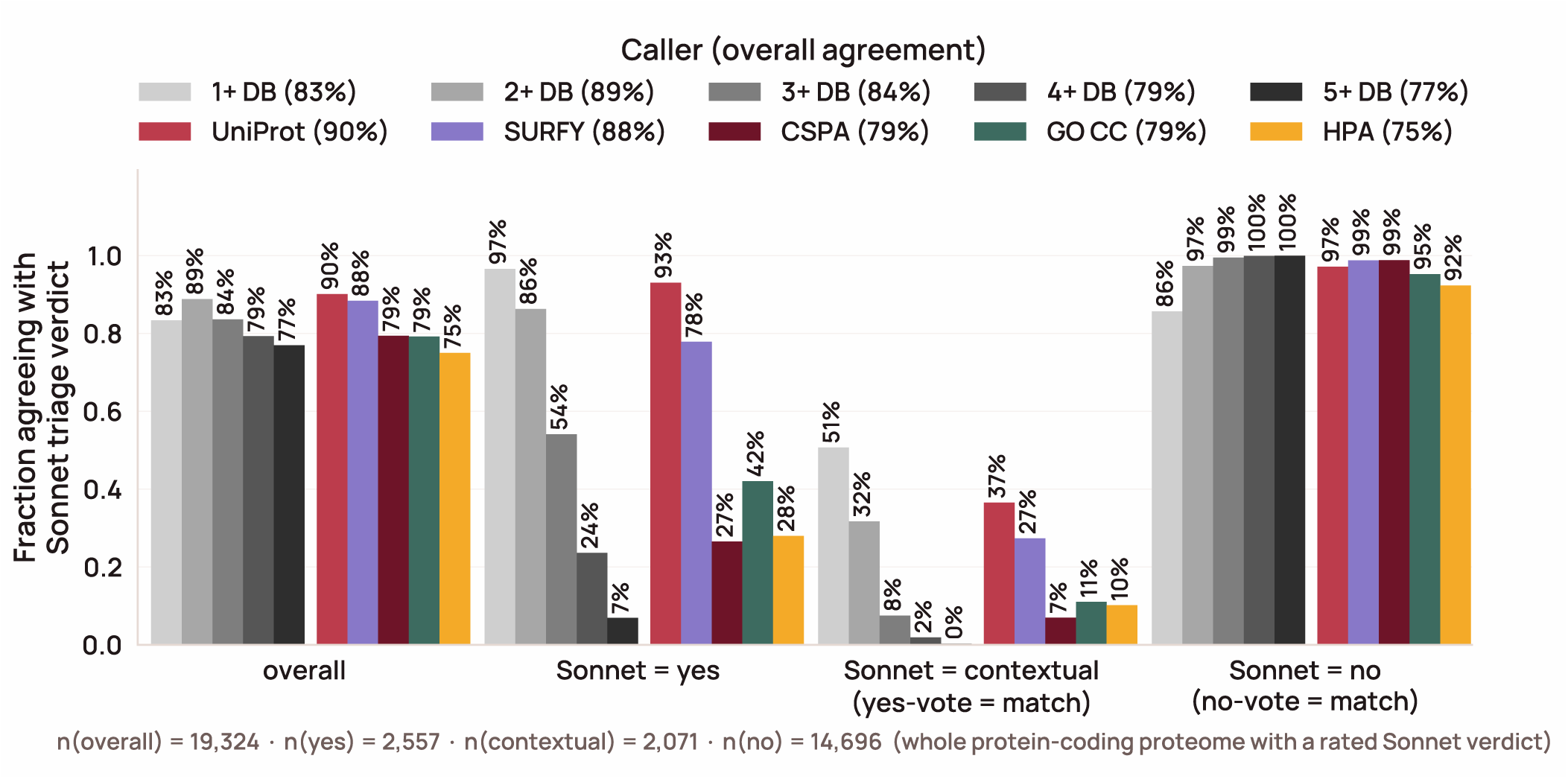
No single surface database recovers more than a fraction of the genome-wide accessible surfaceome the triage agent identifies. Because no hand-curated truth exists at genome scale, agreement is measured across all 19,324 protein-coding genes against the production two-stage triage verdict (yes, contextual, or no) as the reference. A database surface call agrees with an agent yes or contextual; a non-surface call agrees only with no. The databases fall hardest on contextually surface proteins, for which none has an explicit category. [Figure data and script](https://gist.github.com/beccajcarlson/1265c867a3bbb08efd81262789e1f013).

**Supplementary Figure 7.**
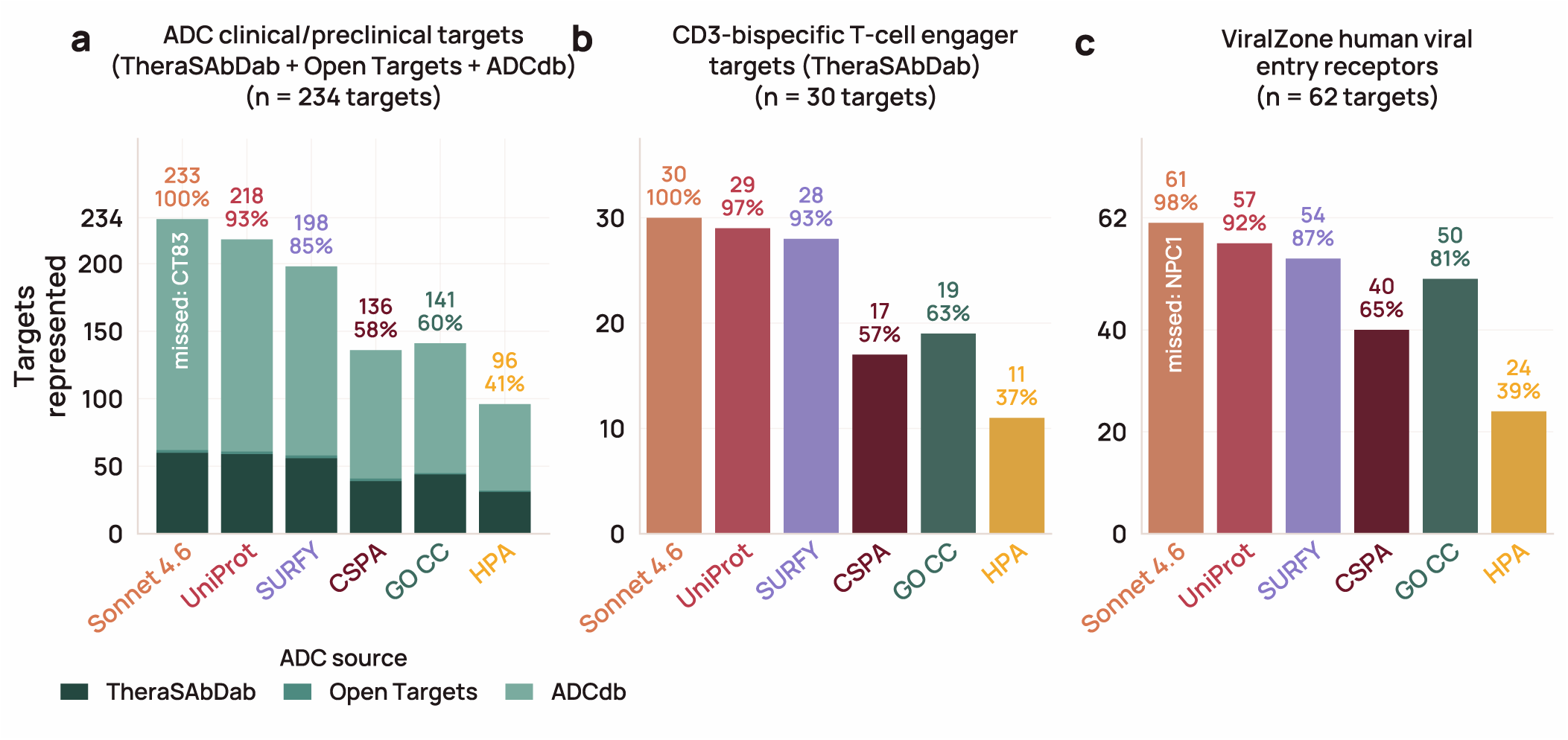
The two-stage Sonnet 4.6 triage agent recovers near-complete coverage of validated surface targets, including targets no individual database calls, across three orthogonal positive-control sets. Each bar is the fraction of an independently curated target set that a source labels surface: the five gating databases (UniProt and CSPA on their SurfaceBench-optimized cutoffs; GO CC, SURFY, and HPA on their native cutoffs) and the two-stage Sonnet 4.6 triage agent (yes or contextual). a) Antibody–drug conjugate targets (n = 234): the union of TheraSAbDab ADCs, Open Targets 26.06 ADC mechanism-of-action targets, and ADCdb antigens, HGNC-resolved, after removing non-cytotoxic conjugates (tedromer/ celmoleukin/cafraglutide), Open Targets family-expansion artifacts, and ADCdb’s secreted/intracellular antigens. b) T-cell engager targets (n = 30): TheraSAbDab CD3-binding bispecifics. c) ViralZone human viral-entry receptors (n = 62). Sonnet covers 233/234 (99.6%) of ADC targets, 30/30 (100%) of TCE targets, and 61/62 (98.4%) of viral-entry receptors, exceeding the best single database (UniProt) in every panel. Eight ADC targets and one TCE target sit in zero of the five databases yet are recovered by Sonnet (ASPH, FMOD, GSDME, HSPA1A, KLK2, KLK3, LRG1, MMP9; KLK2 is in both sets). Most reflect contextual-surface biology (stress-induced chaperones, pyroptotic gasdermin, shed/contextually surface proteases) that the databases’ steady-state definitions miss; two, FMOD and KLK3, are secreted or matrix proteins rather than bona fide surface antigens, and the deep dive called them non-surface. [Figure data and script](https://gist.github.com/beccajcarlson/3ab5df749b576912959c75fe7013d78c).

**Supplementary Figure 8.**
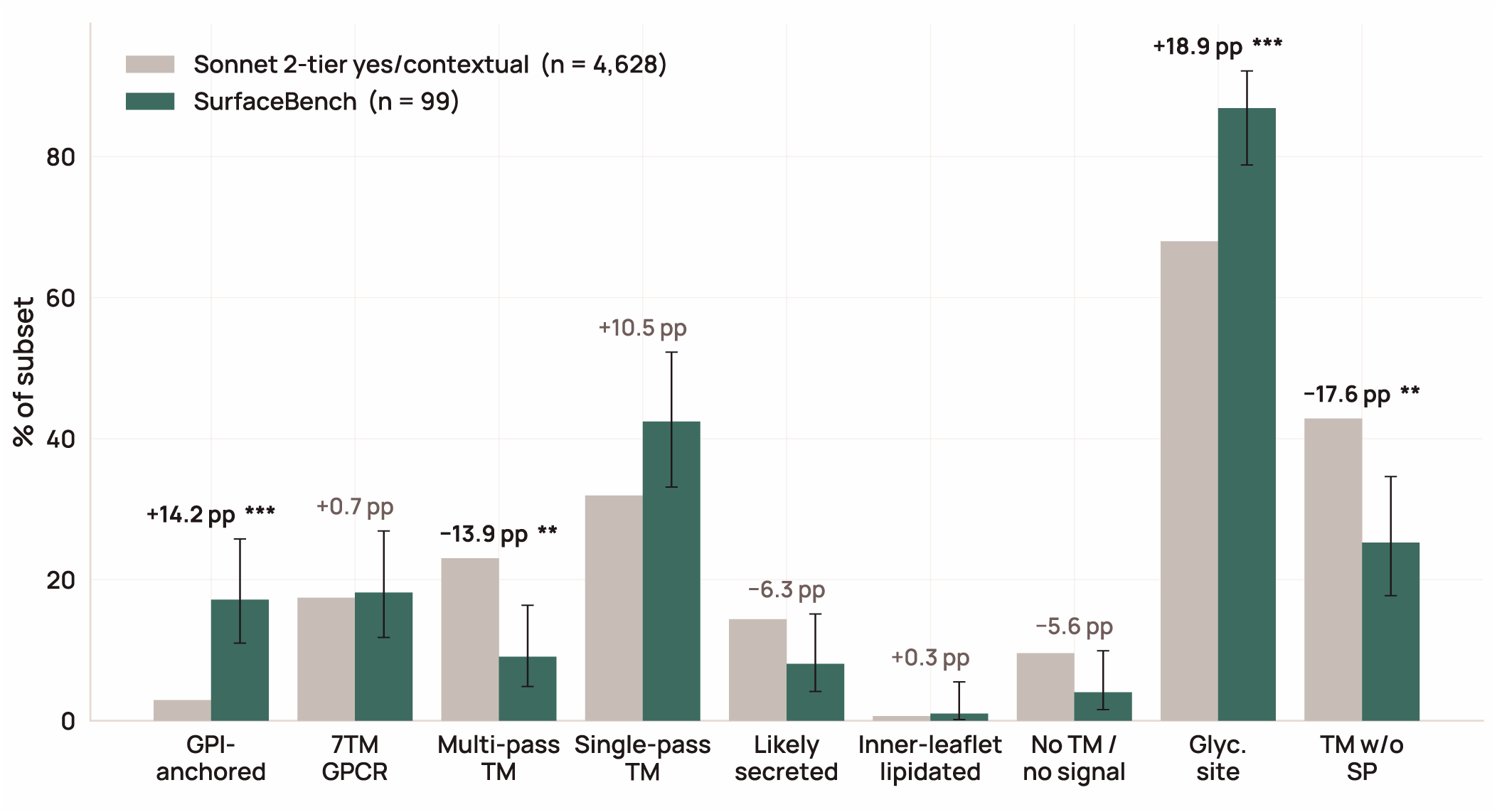
SurfaceBench over-samples the membrane topologies where databases disagree, so its measured accuracy is not a representative estimate of full-catalog performance. The per-class composition of the hand-curated benchmark is compared against the 4,628 proteins the two-stage triage agent calls surface across nine membrane-topology classes, with each class tested by an exact two-tailed binomial test of the benchmark count against the population proportion, Bonferroni-corrected across the nine classes. The benchmark is significantly enriched for glycosylation-site-bearing and GPI-anchored proteins and depleted for multi-pass transmembrane proteins and transmembrane proteins without a signal peptide. [Figure data and script](https://gist.github.com/beccajcarlson/f724dd328f48566a354f0294109a7337).

**Supplementary Figure 9.**
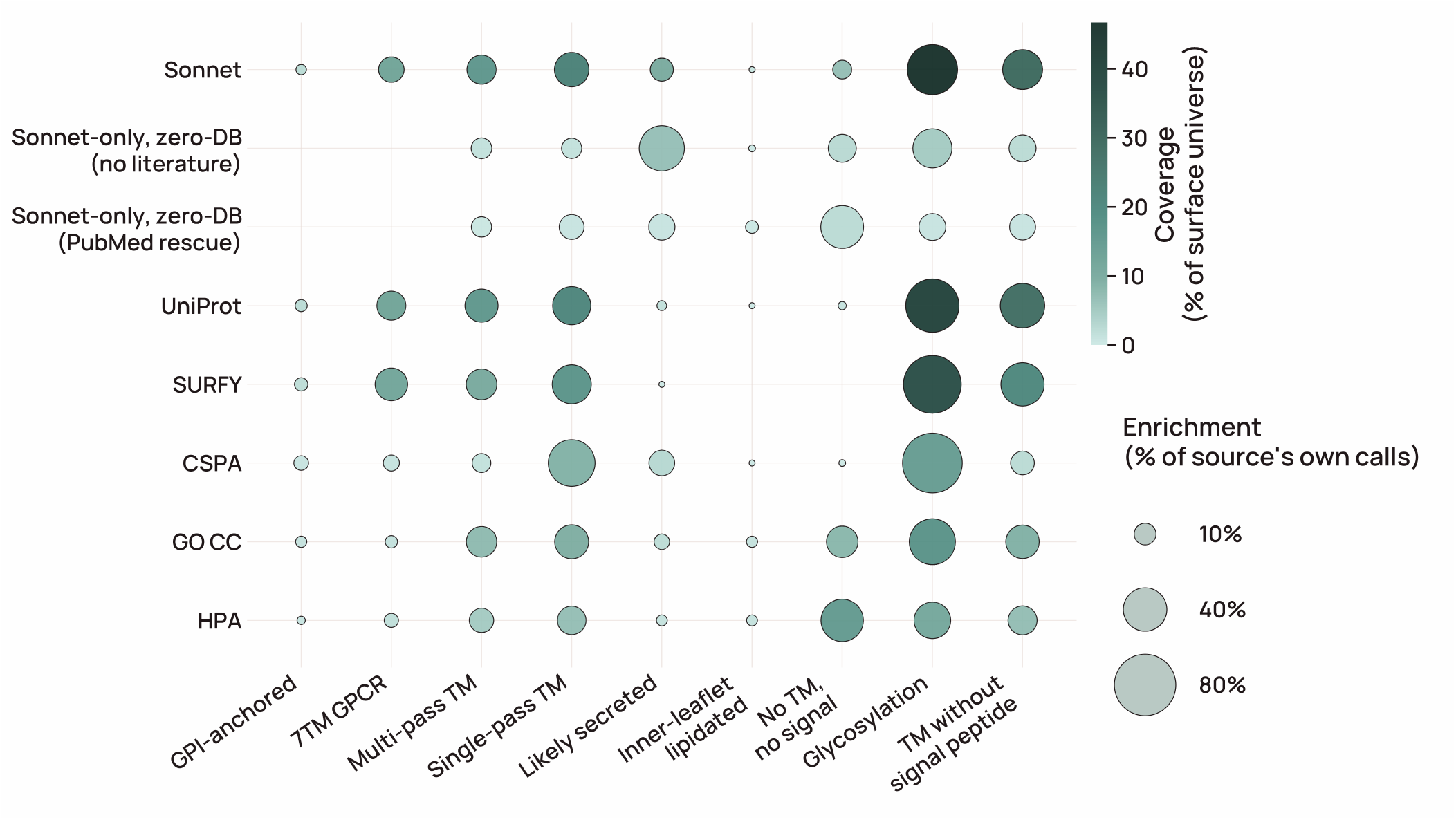
Each surface source is enriched for a distinct set of membrane-topology architectures, but that enrichment and its genome-wide coverage are separate quantities. Bubble matrix of inclusion sources (rows) × topology features (columns) over the candidate-surfaceome universe: dot area is within-source enrichment (fraction of that source’s own calls carrying the feature) and dot color is coverage (fraction of the whole surface universe those proteins represent). CSPA is strongly glycosylation-enriched (∼96% of its own calls) yet covers only ∼14% of the universe, reflecting its N-glycocapture chemistry and cell-line detection limits. SURFY and UniProt show the same glycosylation enrichment at larger scale. Zero-database Sonnet rescues differ by how they were called: those called from the gene summary alone (no literature; 794) are mostly predicted secreted (55%) and glycosylated (41%), whereas those rescued only once PubMed literature was added (314) mostly lack both a transmembrane segment and a signal peptide (49%), so their surface localization is evident only from the literature. [Figure data and script](https://gist.github.com/beccajcarlson/95b0f4cdcaf6a6b91f57539cd1515a25).

**Supplementary Figure 10.**
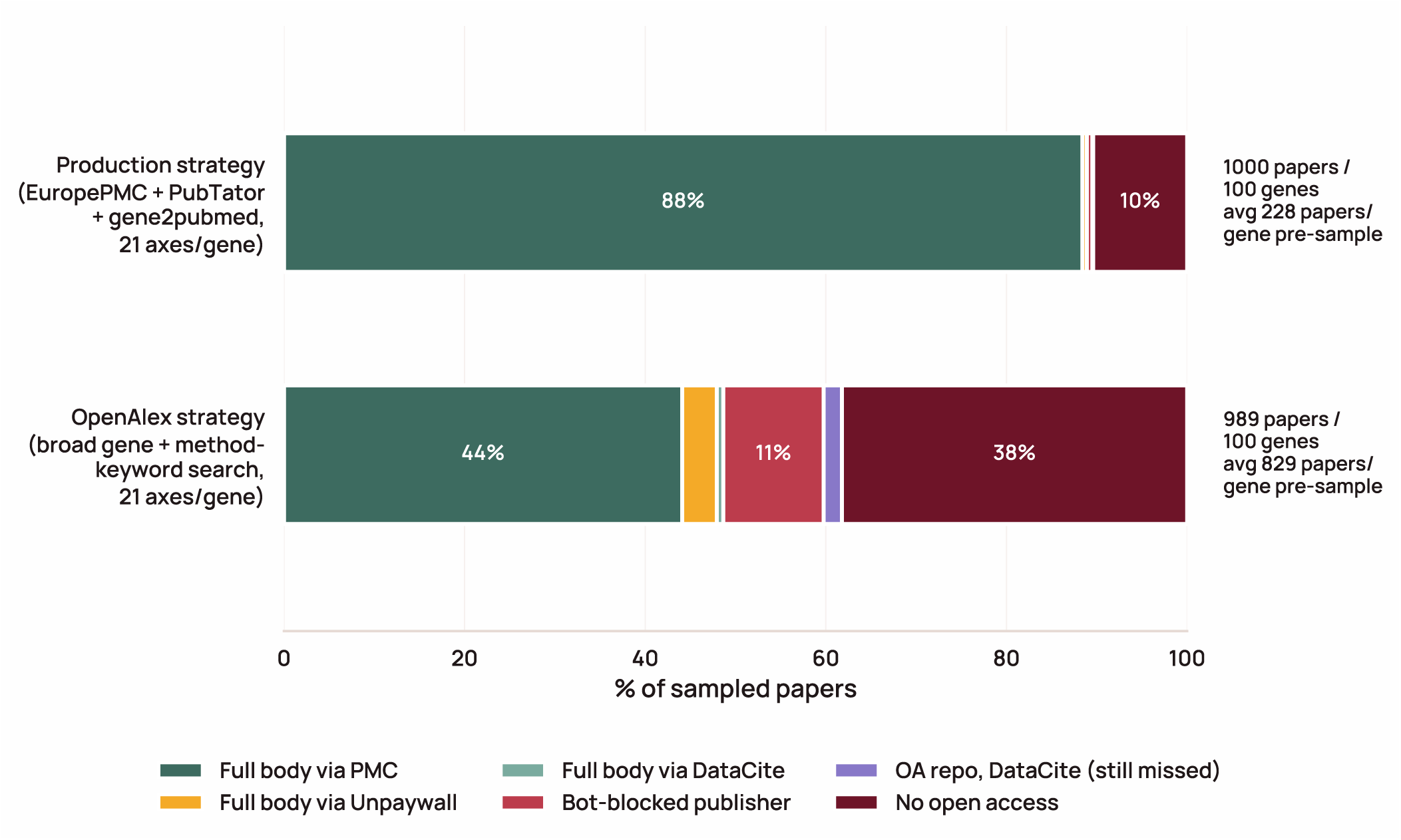
Casting a wider literature net surfaces several times more papers per protein, but most of the additional papers are unreachable, which is why the broader strategy was not adopted. Two discovery strategies were compared on the same random genes and retrieval chain: the production strategy, which searches Europe PMC and adds machine-annotated PubTator mentions, NCBI’s curated gene-to-publication links, and standing topic queries; and a broad alternative issuing one gene-name query plus method-keyword queries against the OpenAlex index, which also reaches preprints and non-PubMed journals. OpenAlex discovered about 3.6-fold more papers per protein, but the reachable full-text fraction fell from 89% to 49%, and the extra papers were mostly inaccessible or off-topic (Supplementary Note 1). [Figure data and script](https://gist.github.com/beccajcarlson/cbc950dad1c3a6595fd5018cdb6b030d).

**Supplementary Figure 11.**
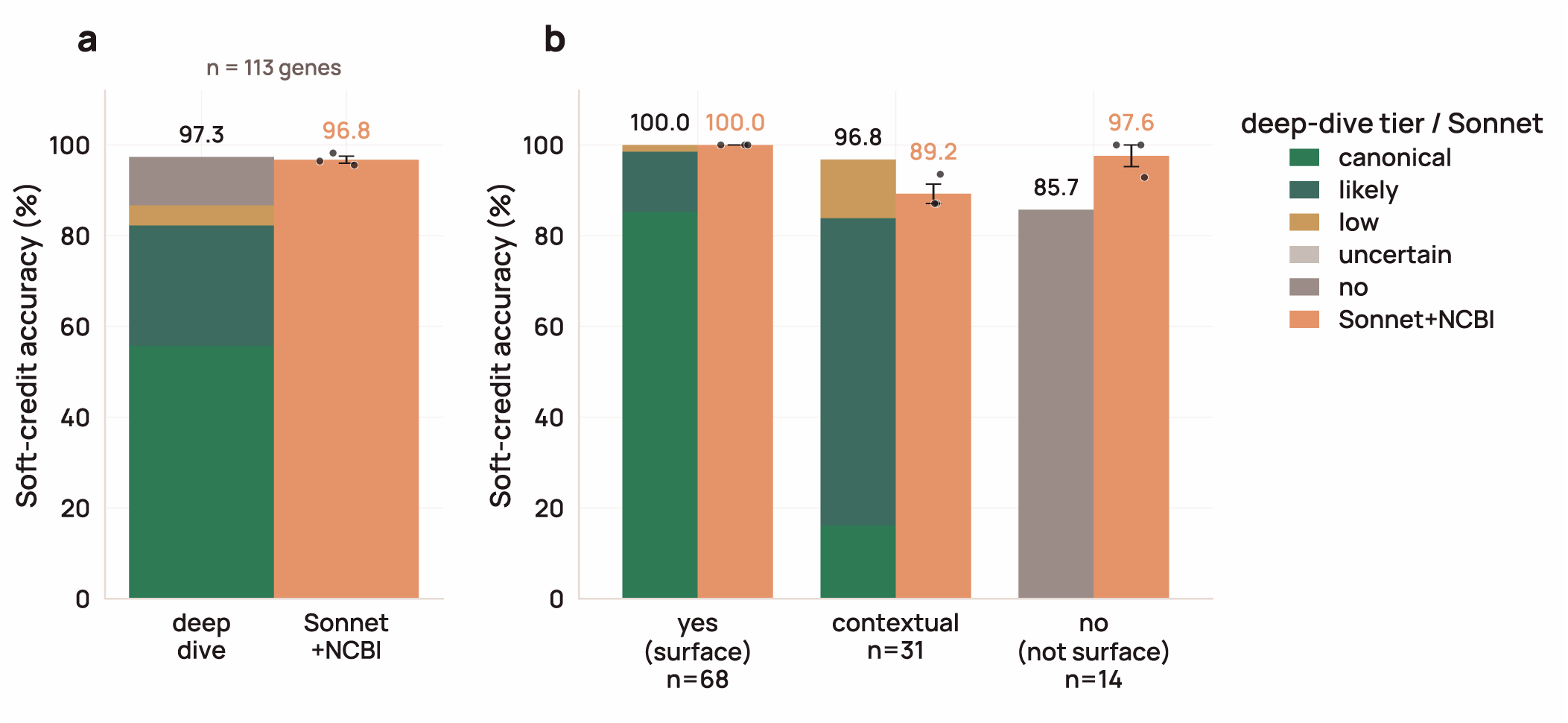
For SurfaceBench proteins subjected to deep-dive analysis, the evidence-based assessment matched the soft-credit accuracy of Sonnet+NCBI triage, while the surface tiers reflected the difficulty of each case. For the 113 deep-dived SurfaceBench genes, deep-dive and Sonnet+NCBI triage calls are scored against the curated verdict under the soft-credit rule. a) Overall soft-credit accuracy. b) Accuracy split by ground-truth bucket (yes / contextual / no). Deep-dive bars are stacked by the tier assigned to correctly classified genes (palette as in Figure 5a): ground-truth-yes genes are called mostly canonical (58 of 68), contextual genes mostly likely (21 of 30 correct), and the correct non-surface calls all land in no. Sonnet+NCBI bars are the mean of three replicates (dots; error bar SEM); the deep dive was run once, so its bar has no spread. The two methods agree overall (97.3% vs 96.8%) and on ground-truth-yes genes (both 100%); the deep dive scores higher on contextual genes (96.8% vs 89.2%) and lower on non-surface genes (85.7% vs 97.6%; both misses were called low). None of these differences is significant by an exact McNemar test on paired gene-level correctness (overall p = 1.0; contextual p = 0.63; non-surface p = 0.50). [Figure data and script](https://gist.github.com/beccajcarlson/d88f7c7a135bf222ccc5883b9c1a2eb4).

**Supplementary Figure 12.**
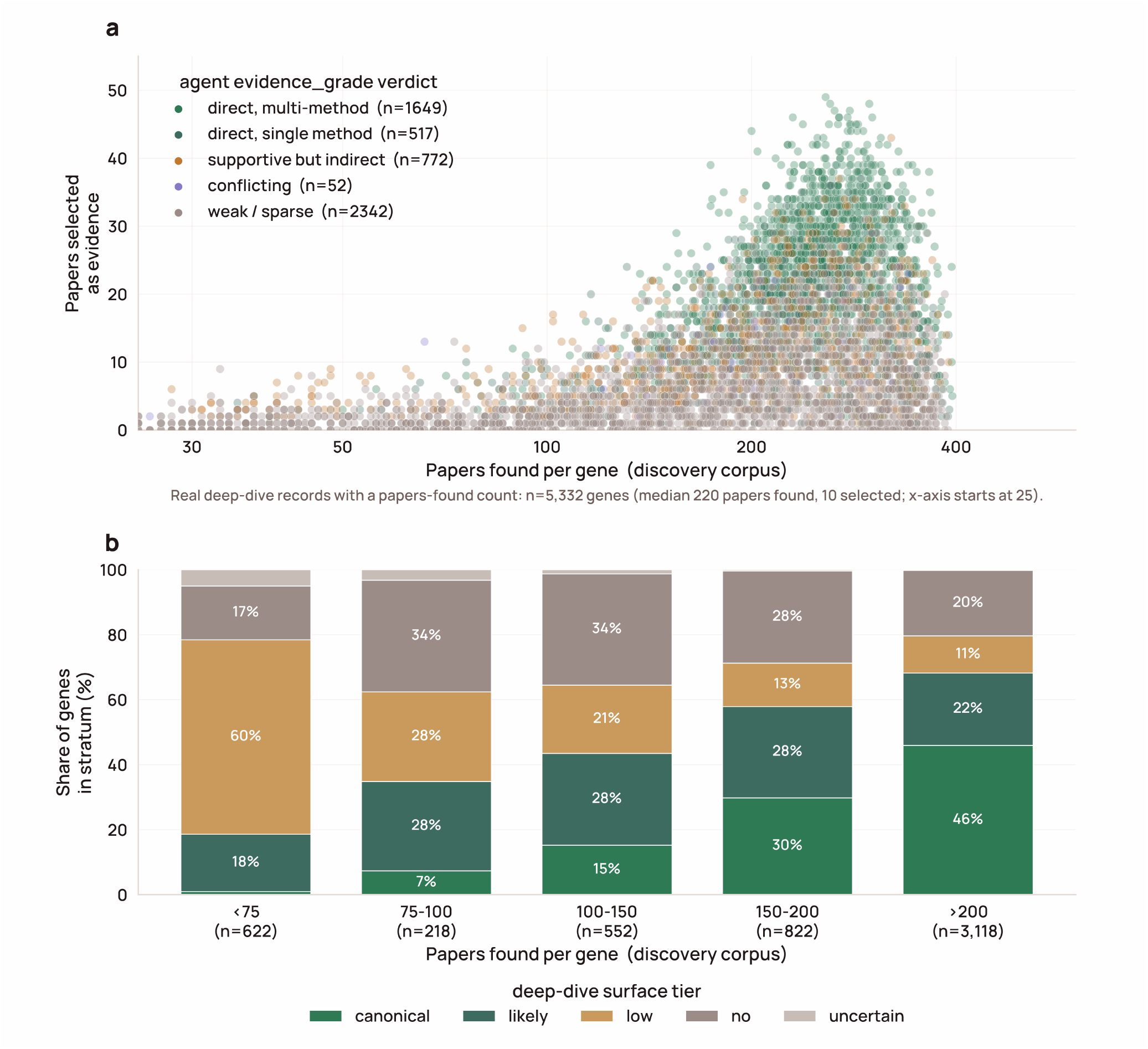
The number of papers retained as evidence per protein rises with the size of the discovery corpus, and the agent’s evidence-quality grade tracks the depth of the retained set. a) For each deep-dived protein (n = 5,332), the number of papers found during literature discovery (horizontal axis, log scale) against the number kept as evidence after selection (vertical axis, linear scale); median 220 found and 10 kept. Points are colored by the agent’s evidence-quality grade: direct (multiple independent assays), direct (single assay), supportive but indirect, conflicting, or weak or sparse. b) Deep-dive surface tier of proteins grouped by the number of papers found. The share called canonical rises from 1% of proteins with fewer than 75 papers to 46% of those with 200 or more, while the share called low falls from 60% to 11%. Proteins with fewer than 100 papers found that were called surface by UniProt are flagged on the viewer, as possible surface proteins with too little literature to classify. [Figure data and script](https://gist.github.com/beccajcarlson/cb550702344d7626ee80804bc7f6c549).

**Supplementary Figure 13.**
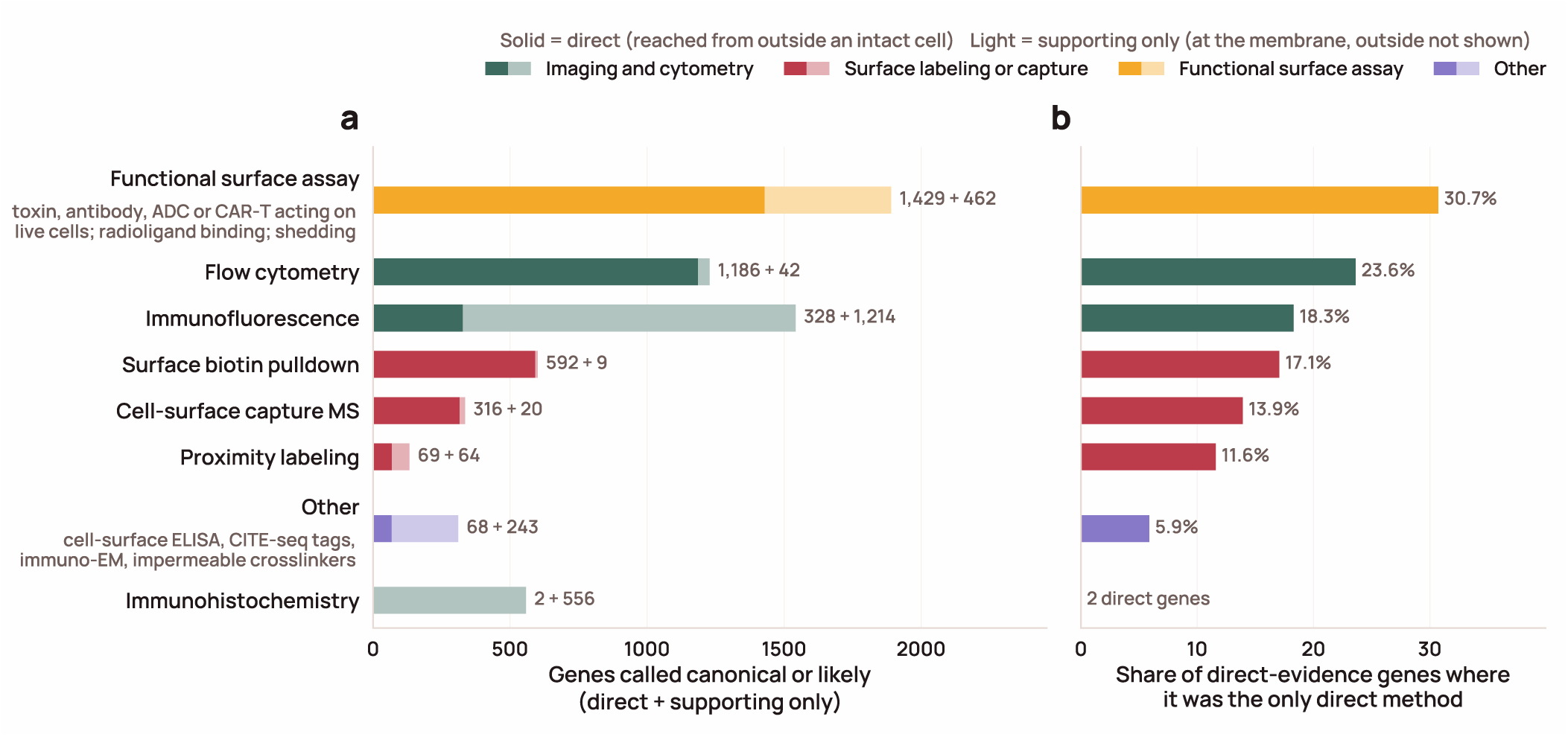
Functional assays and flow cytometry provide most of the direct surface evidence, while immunofluorescence and immunohistochemistry mostly place proteins at the membrane without showing they are reachable from outside. For the 3,035 deep-dived proteins called canonical or likely, each experiment the agent extracted from the cited literature is classified by method and judged direct (an antibody or probe reached the protein from outside an intact cell: live or non-permeabilized staining of an extracellular epitope or tag, membrane-impermeable labeling, or a functional readout that requires surface access) or supporting (places the protein at the plasma membrane without showing its outside is reachable: membrane-rim staining, fluorescent-protein fusions, or membranous immunohistochemistry). a) Proteins with direct evidence from each method (solid) and proteins for which that method reached only the supporting level (light). b) Share of the 2,203 proteins with any direct evidence for which the method was the only direct method; immunohistochemistry had two direct proteins, too few for a share. Colors group methods into imaging and cytometry, surface labeling or capture, functional surface assays, and other methods (for example, cell-surface ELISA, CITE-seq antibody tags, immuno-electron microscopy on intact cells, and membrane-impermeable crosslinking). Functional surface assays include toxin, antibody, ADC, or CAR-T activity against live target-positive cells, live-cell radioligand binding, ectodomain shedding, and extracellular substrate transport. [Figure data and script](https://gist.github.com/beccajcarlson/141184baa53bc889d2c08692b931982b).

**Supplementary Figure 14.**
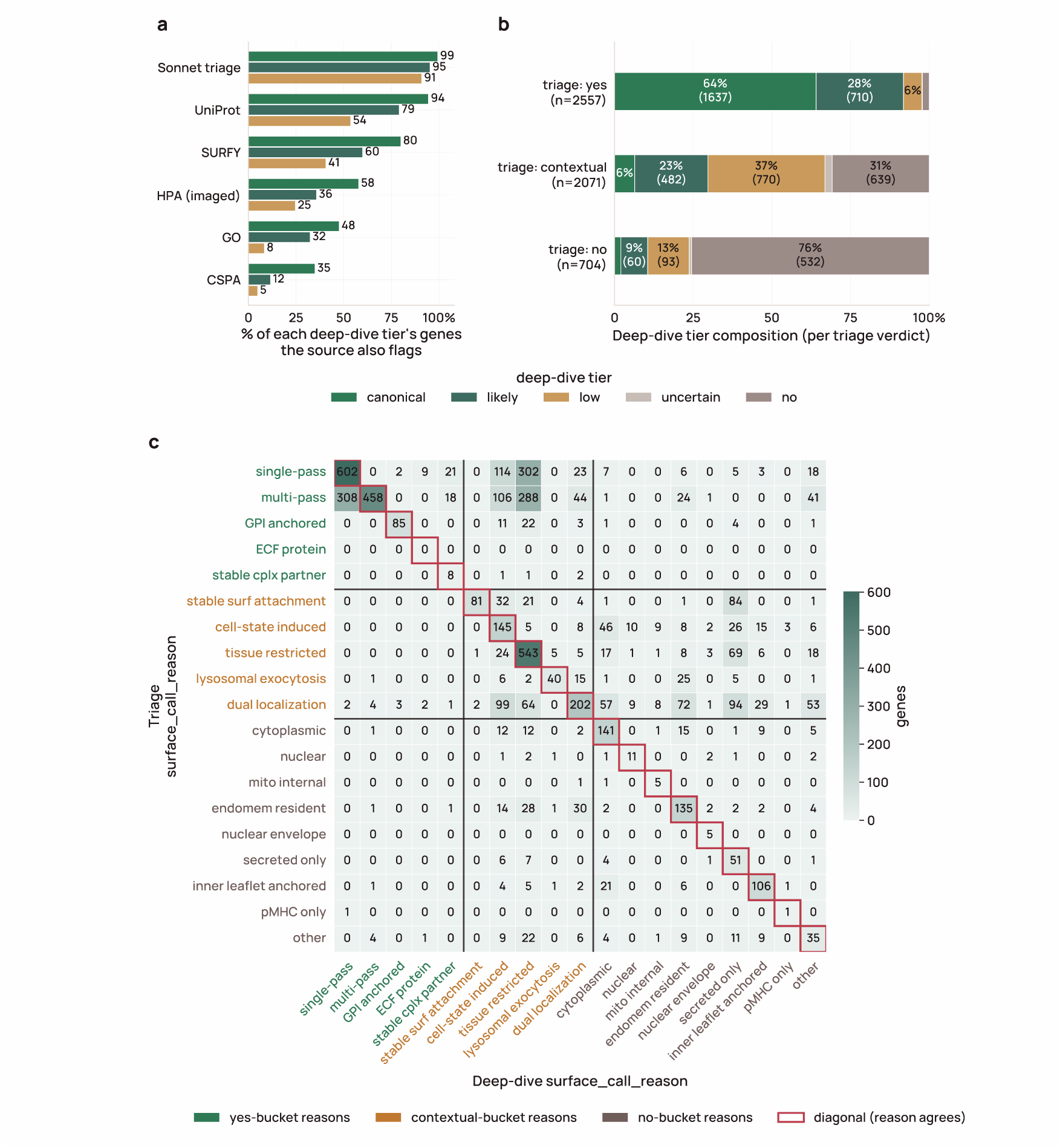
Triage and database calls agree with full-text deep-dive curation most strongly for high-confidence surface proteins (n = 5,332 deep-dived proteins). a) Fraction of deep-dive canonical, likely, and low-tier proteins also flagged as surface by each database and by the triage agent; each fraction is taken over all proteins in that tier, so a protein absent from a database counts as not flagged, except for HPA, which is scored only over the 2,917 proteins it imaged because absence from HPA reflects a missing antibody rather than a negative call. b) Resolution of each triage verdict across the five deep-dive tiers. Triage-yes is 64% canonical and 28% likely; triage-contextual spreads across likely, low, and no; triage-no remains 76% no, with 22% upgraded to likely or low by the deep-dive pipeline. c) Triage reason (rows) versus deep-dive structured reason (columns); exact agreement is 50%. [Figure data and script](https://gist.github.com/beccajcarlson/152a4d8b4a1d156e2afaa73d205294b6).

**Supplementary Figure 15.**
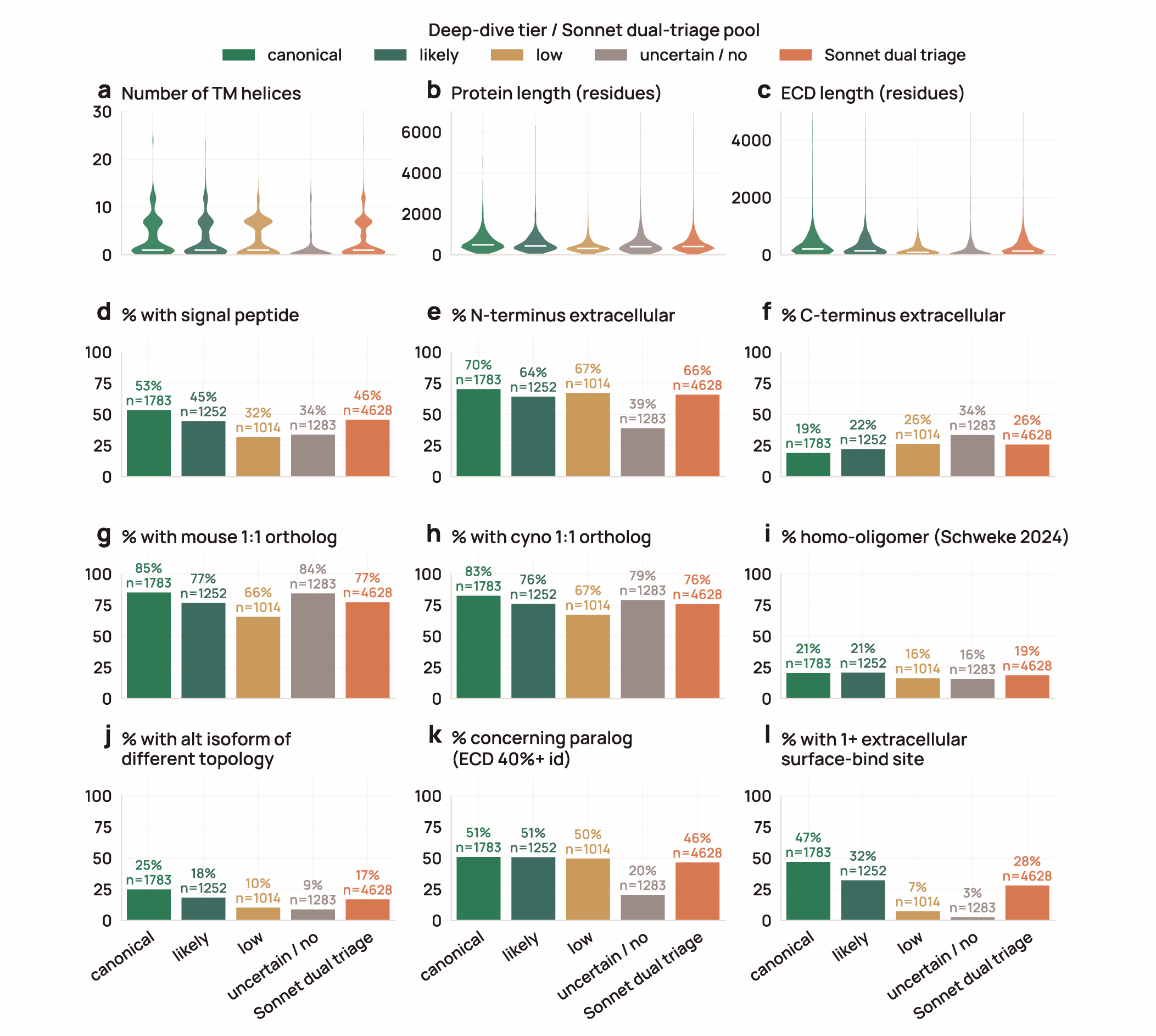
Precomputed deterministic features distribute as expected for surface proteins across deep-dive tiers and the triage-positive pool. Twelve topology and structural features (TM-helix count, protein length and ECD length, signal peptide, N-and C-terminal orientation, mouse and cynomolgus 1:1 orthology, homo-oligomerization, alt-isoform topology change, concerning paralog, and extracellular SURFACE-Bind site) compared across the canonical, likely, low, and uncertain/ non-surface tiers and the triage-positive pool: all 4,628 proteins the two-stage Sonnet 4.6 triage called yes or contextual (all have a canonical DeepTMHMM topology), deep-dived or not, with features taken from the genome-wide tables. [Figure data and script](https://gist.github.com/beccajcarlson/57cf3cc3db903ab39bc0ba315ce5e4d5).

**Supplementary Figure 16.**
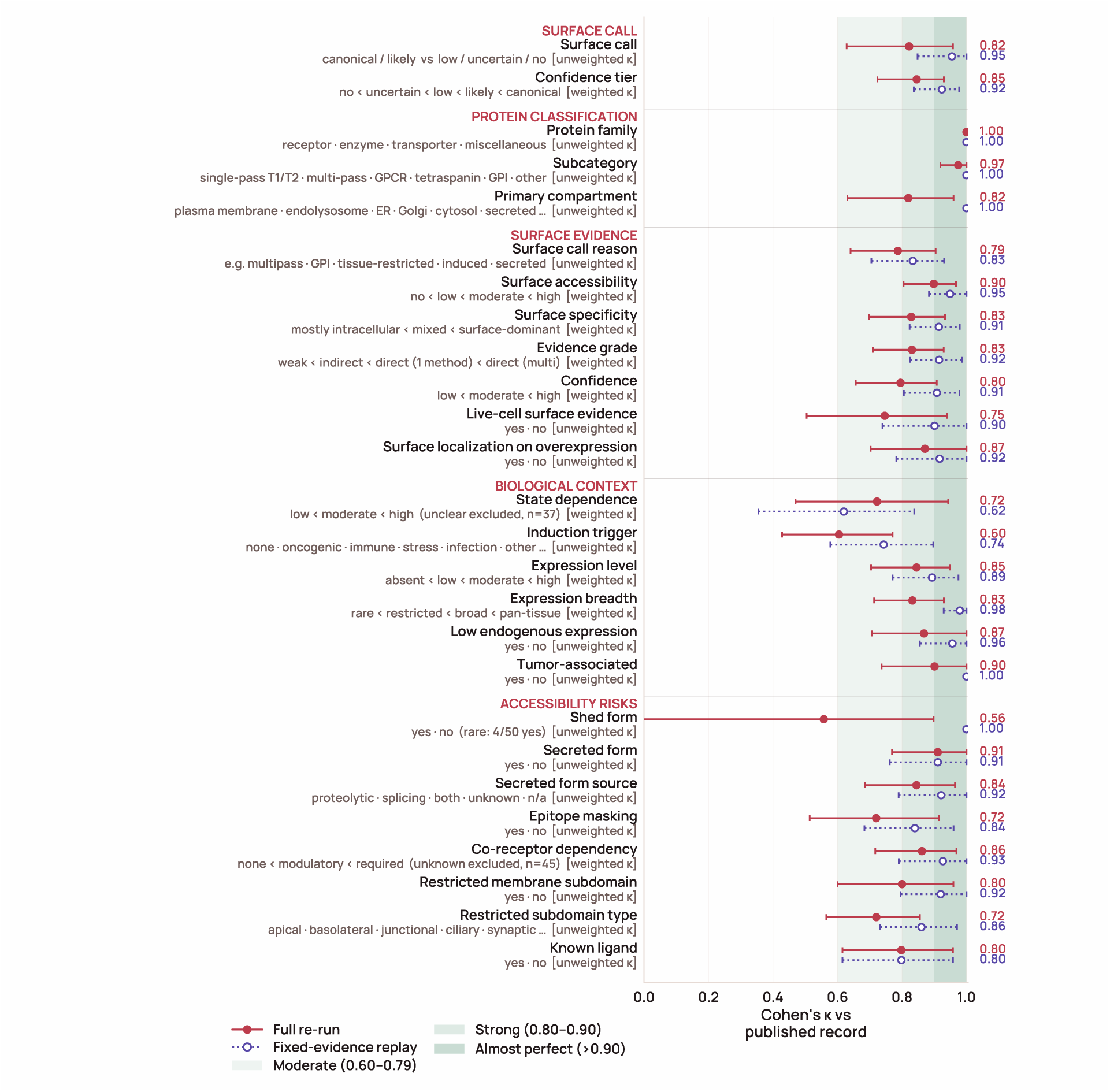
Deep-dive records are reproducible, and most run-to-run differences come from which papers are retrieved. For 50 random deep-dive proteins, we compared each of the 24 LLM-derived fields, plus the surface call and tier, with the published record under two re-analyses. The full re-run (maroon, solid) repeated the pipeline from literature discovery onward; the fixed-evidence replay (lavender, dotted) reused each record’s selected evidence and re-ran only the section builders and synthesizer. Points show Cohen’s κ versus the published record, with 95% intervals from 5,000 bootstrap resamples of proteins. Ordered fields use linearly weighted κ. State dependence and co-receptor dependency exclude unclear and unknown calls (n shown). Green shading marks moderate (0.60–0.79), strong (0.80–0.90), and almost perfect (>0.90) agreement (McHugh, 2012). Induction trigger is scored as a single dominant trigger, although several can apply to one protein. Shed form is rare (4 of 50 proteins), which lowers κ despite 92% raw agreement. [Figure data and script](https://gist.github.com/beccajcarlson/14eec28d77e0d529213815c6af4283e0).

### SUPPLEMENTARY NOTE 1. LIMITATIONS OF FULL-TEX T RETRIEVAL BY AGENTS

A major limitation of this work is that much of the relevant literature is not freely accessible to researchers or automated agents. Full text, not just abstracts, matters for such pipelines, as shown previously for protein subcellular associations and other features (Westergaard et al., 2018). On a random sample of the discovery corpus (100 proteins, 10 papers each), 89% of candidate papers were recoverable as full text (Figure S10). On the same cohort, a broader OpenAlex search returned 3.6-fold more papers per protein (mean 829 vs 228) but only 49% were bot-reachable, versus 89% for our targeted queries. Of the ∼33 incremental papers Unpaywall could recover across the 100 proteins, 16 of 20 manually audited contained the protein name in passing only. The reachable, on-topic gain was negligible, and OpenAlex’s free API, whose daily rate limit we exhausted even at audit scale, would have required a paid tier for a proteome-wide sweep, so we did not adopt it.

The bot-accessible corpus was effectively PMC (88% of a random sample of papers), whose content is heavily weighted toward NIH-and partner-funder-mandated deposits — a substantial narrowing of the global research record. Repositories like bioRxiv were nominally open but unreachable by per-article HTTP retrieval (full-text data mining access is delivered only as monthly bulk dumps via a requester-pays S3 bucket). Zenodo and arXiv are bot-accessible, but their DOIs register through DataCite, where Unpaywall’s coverage is sparser than for Crossref (DataCite DOIs were only 2.7% of our sample), and even a dedicated DataCite fallback reached few of them. The obstacle is the metadata, not the PDF: PDF parsing is imperfect but well supported, whereas DataCite metadata is created across many repositories without a shared standard for where the full text lives; in our sample, not one record populated the schema’s optional content-URL field. We instead scraped the landing page’s “citation_ pdf_url” tag, which succeeded for only seven records: four from arXiv, whose fixed URL scheme we parse directly, and three elsewhere — one Zenodo and two institutional repositories. What we missed was mostly repository theses and off-topic records rather than research papers. No current tool simultaneously maximizes visibility to the research community and to automated agents.

To partially close this gap, we deposited this paper to bioRxiv (for community visibility and a Crossref-registered DOI) and to Zenodo as both a PDF and a machine-readable text file (since Unpaywall linking surfaces direct full-text URLs), with the Zenodo record linked to the bioRxiv DOI so that agents can retrieve it whichever registry a tool indexes. Broader adoption of dual deposition, or a cross-registry resolver, would expand the corpus available to literature-aware LLM pipelines like ours.

Finally, the accuracy of these and all other LLM-generated calls ultimately depends on the reliability of the claims in the machine-readable corpus. Published assertions do not always reflect the underlying data: a claim about protein localization may rely on a poorly validated antibody or binding reagent, arise from an artificial experimental system, or lack adequate support from the reported data. The literature is also enriched for positive findings and depleted of negative or null results, though such findings would broaden what models learn from. We therefore need both to publish results that do not support the original hypothesis (including null findings and evidence for alternative hypotheses) and to make post-publication critiques openly available. Journal-club discussions, for example, could be recorded, summarized with AI, and released, adding scientific discourse to the machine-readable record. As LLMs become widely accessible and increasingly serve as a starting point for scientific reasoning, a public record of post-publication review will help ensure that models learn not only from published claims, but also from the evidence and critical reasoning needed to evaluate them.

## REFERENCES

Balbi, P.E.M., Sadek, A., Marchand, A., Yu, T.-Y., Damjanovic, J., Georgeon, S., Schmidt, J., Fulle, S., Yang, C., Khakzad, H., Correia, B.E., 2026. Mapping targetable sites on the human surfaceome for the design of novel binders. Proc. Natl. Acad. Sci. 123, e2506269123. 10.1073/pnas.2506269123

Bausch-Fluck, D., Goldmann, U., Müller, S., van Oostrum, M., Müller, M., Schubert, O.T., Wollscheid, B., 2018. The in silico human surfaceome. Proc. Natl. Acad. Sci. 115, E10988–E10997. 10.1073/pnas.1808790115

Bausch-Fluck, D., Hofmann, A., Bock, T., Frei, A.P., Cerciello, F., Jacobs, A., Moest, H., Omasits, U., Gundry, R.L., Yoon, C., Schiess, R., Schmidt, A., Mirkowska, P., Härtlová, A., Van Eyk, J.E., Bourquin, J.-P., Aebersold, R., Boheler, K.R., Zandstra, P., Wollscheid, B., 2015. A Mass Spectrometric-Derived Cell Surface Protein Atlas. PLoS ONE 10, e0121314. 10.1371/journal.pone.0121314

Bertoni, D., Tsenkov, M., Magana, P., Nair, S., Pidruchna, I., Querino Lima Afonso, M., Midlik, A., Paramval, U., Lawal, D., Tanweer, A., Last, M., Patel, R., Laydon, A., Lasecki, D., Dietrich, N., Tomlinson, H., Žídek, A., Green, T., Kovalevskiy, O., Lau, A., Kandathil, S., Bordin, N., Sillitoe, I., Mirdita, M., Jones, D., Orengo, C., Steinegger, M., Fleming, J.R., Velankar, S., 2026. AlphaFold Protein Structure Database 2025: a redesigned interface and updated structural coverage. Nucleic Acids Res. 54, D358–D362. 10.1093/nar/gkaf1226

Betts, M.R., Brenchley, J.M., Price, D.A., De Rosa, S.C., Douek, D.C., Roederer, M., Koup, R.A., 2003. Sensitive and viable identification of antigen-specific CD8+ T cells by a flow cytometric assay for degranulation. J. Immunol. Methods 281, 65–78. 10.1016/s0022-1759(03)00265-5

Brase, J., 2009. DataCite -A Global Registration Agency for Research Data, in: Proceedings of the 2009 Fourth International Conference on Cooperation and Promotion of Information Resources in Science and Technology, COINFO ’09. IEEE Computer Society, USA, pp. 257–261. 10.1109/COINFO.2009.66

Buniello, A., Suveges, D., Cruz-Castillo, C., Llinares, M.B., Cornu, H., Lopez, I., Tsukanov, K., Roldán-Romero, J.M., Mehta, C., Fumis, L., McNeill, G., Hayhurst, J.D., Martinez Osorio, R.E., Barkhordari, E., Ferrer, J., Carmona, M., Uniyal, P., Falaguera, M.J., Rusina, P., Smit, I., Schwartzentruber, J., Alegbe, T., Ho, V.W., Considine, D., Ge, X., Szyszkowski, S., Tsepilov, Y., Ghoussaini, M., Dunham, I., Hulcoop, D.G., McDonagh, E.M., Ochoa, D., 2025. Open Targets Platform: facilitating therapeutic hypotheses building in drug discovery. Nucleic Acids Res. 53, D1467–D1475. 10.1093/nar/gkae1128

Camon, E.B., Barrell, D.G., Dimmer, E.C., Lee, V., Magrane, M., Maslen, J., Binns, D., Apweiler, R., 2005. An evaluation of GO annotation retrieval for BioCreAtIvE and GOA. BMC Bioinformatics 6 Suppl 1, S17. 10.1186/1471-2105-6-S1-S17

Carette, J.E., Raaben, M., Wong, A.C., Herbert, A.S., Obernosterer, G., Mulherkar, N., Kuehne, A.I., Kranzusch, P.J., Griffin, A.M., Ruthel, G., Cin, P.D., Dye, J.M., Whelan, S.P., Chandran, K., Brummelkamp, T.R., 2011. Ebola virus entry requires the cholesterol transporter Niemann-Pick C1. Nature 477, 340–343. 10.1038/nature10348

Carlson, R.J., Smallegan, M.J., 2026a. The accessible human surfaceome. 10.5281/zenodo.23445521

Carlson, R.J., Smallegan, M.J., 2026b. accessible-surfaceome: an LLM-curated atlas of the accessible human surfaceome. 10.5281/zenodo.23455406

Chen, E.Y., Tan, C.M., Kou, Y., Duan, Q., Wang, Z., Meirelles, G.V., Clark, N.R., Ma’ayan, A., 2013. Enrichr: interactive and collaborative HTML5 gene list enrichment analysis tool. BMC Bioinformatics 14, 128. 10.1186/1471-2105-14-128

Chi, K.N., Gedye, C., Belkoff, L., Barata, P.C., Bernard-Tessier, A., Graff, J.N., Matsubara, N., Sandhu, S., Schweizer, M.T., Stein, M.N., Von Amsberg, G., Siddiqui, B.A., Wei, X.X., Hosseini, M., Kramer, K., Moussa, F., Wang, S.C.-E., Dibaj, S., Gaut, D.L., Castro, E., 2026. A phase 3 randomized, double-blind, placebo-controlled study of a pasritamig plus best supportive care in metastatic castration-resistant prostate cancer. J. Clin. Oncol. 44, TPS5136–TPS5136. 10.1200/JCO.2026.44.16_suppl.TPS5136

Coyote-Maestas, W., Nedrud, D., He, Y., Schmidt, D., 2022. Determinants of trafficking, conduction, and disease within a K+ channel revealed through multiparametric deep mutational scanning. eLife 11, e76903. 10.7554/eLife.76903

Dana, J.M., Gutmanas, A., Tyagi, N., Qi, G., O’Donovan, C., Martin, M., Velankar, S., 2019. SIFTS: updated Structure Integration with Function, Taxonomy and Sequences resource allows 40-fold increase in coverage of structure-based annotations for proteins. Nucleic Acids Res. 47, D482–D489. 10.1093/nar/gky1114

de Goeij, B.E.C.G., Vink, T., ten Napel, H., Breij, E.C.W., Satijn, D., Wubbolts, R., Miao, D., Parren, P.W.H.I., 2016. Efficient Payload Delivery by a Bispecific Antibody–Drug Conjugate Targeting HER2 and CD63. Mol. Cancer Ther. 15, 2688–2697. 10.1158/1535-7163.MCT-16-0364

Delaveris, C.S., Loudermilk, R.P., Pandey, A., Remesh, S.G., Peters-Clarke, T.M., Ganjave, S.D., Dougherty, W.J.N., Delavan, H.M., Wang, C., Ling, J., Camara Serrano, J.A., Salangsang, F., Ding, C.-K.C., Greenland, N., Chu, C.E., Porten, S., Steri, V., Chou, J., Evans, M.J., Leung, K.K., Wells, J.A., 2026. Autophagolysosomal exocytosis inverts Src kinase onto the cell surface in cancer. Science 391, eaec1778. 10.1126/science.aec1778

Ding, Y., Zuo, Y., Zhang, B., Fan, Y., Xu, G., Cheng, Z., Ma, S., Fang, S., Tian, A., Gao, D., Xu, X., Wang, Q., Jing, Y., Jiang, M., Xiong, M., Li, J., Han, Z., Sun, S., Wang, S., He, F., Yang, J., Qu, J., Zhang, W., Liu, G.-H., 2025. Comprehensive human proteome profiles across a 50-year lifespan reveal aging trajectories and signatures. Cell 188, 5763–5784.e26. 10.1016/j.cell.2025.06.047

Dyer, S.C., Austine-Orimoloye, O., Azov, A.G., Barba, M., Barnes, I., Barrera-Enriquez, V.P., Becker, A., Bennett, R., Beracochea, M., Berry, A., Bhai, J., Bhurji, S.K., Boddu, S., Branco Lins, P.R., Brooks, L., Ramaraju, S.B., Campbell, L.I., Martinez, M.C., Charkhchi, M., Cortes, L.A., Davidson, C., Denni, S., Dodiya, K., Donaldson, S., El Houdaigui, B., El Naboulsi, T., Falola, O., Fatima, R., Genez, T., Martinez, J.G., Gurbich, T., Hardy, M., Hollis, Z., Hunt, T., Kay, M., Kaykala, V., Lemos, D., Lodha, D., Mathlouthi, N., Merino, G.A., Merritt, R., Mirabueno, L.P., Mushtaq, A., Hossain, S.N., Pérez-Silva, J.G., Perry, M., Piližota, I., Poppleton, D., Prosovetskaia, I., Raj, S., Salam, A.I.A., Saraf, S., Saraiva-Agostinho, N., Sinha, S., Sipos, B., Sitnik, V., Steed, E., Suner, M.-M., Surapaneni, L., Sutinen, K., Tricomi, F.F., Tsang, I., Urbina-Gómez, D., Veidenberg, A., Walsh, T.A., Willhoft, N.L., Allen, J., Alvarez-Jarreta, J., Chakiachvili, M., Cheema, J., da Rocha, J.B., De Silva, N.H., Giorgetti, S., Haggerty, L., Ilsley, G.R., Keatley, J., Loveland, J.E., Moore, B., Mudge, J.M., Naamati, G., Tate, J., Trevanion, S.J., Winterbottom, A., Flint, B., Frankish, A., Hunt, S.E., Finn, R.D., Freeberg, M.A., Harrison, P.W., Martin, F.J., Yates, A.D., 2025. Ensembl 2025. Nucleic Acids Res. 53, D948–D957. 10.1093/nar/gkae1071

Edfors, F., Hober, A., Linderbäck, K., Maddalo, G., Azimi, A., Sivertsson, Å., Tegel, H., Hober, S., Szigyarto, C.A.-K., Fagerberg, L., von Feilitzen, K., Oksvold, P., Lindskog, C., Forsström, B., Uhlen, M., 2018. Enhanced validation of antibodies for research applications. Nat. Commun. 9, 4130. 10.1038/ s41467-018-06642-y

Fang, H., Jiang, L., Leprevost, F. da V., Jian, R., Chan, J., Glinos, D., Lappalainen, T., Nesvizhskii, A.I., Reiner, A.P., Consortium, Gte., Snyder, M.P., Tang, H., 2025. Regulation of protein abundance in normal human tissues. 10.1101/2025.01.10.25320181

Ferguson, C., Araújo, D., Faulk, L., Gou, Y., Hamelers, A., Huang, Z., Ide-Smith, M., Levchenko, M., Marinos, N., Nambiar, R., Nassar, M., Parkin, M., Pi, X., Rahman, F., Rogers, F., Roochun, Y., Saha, S., Selim, M., Shafique, Z., Sharma, S., Stephenson, D., Talo’, F., Thouvenin, A., Tirunagari, S., Vartak, V., Venkatesan, A., Yang, X., McEntyre, J., 2021. Europe PMC in 2020. Nucleic Acids Res. 49, D1507–D1514. 10.1093/nar/gkaa994

Gene Ontology Consortium, 2026. The Gene Ontology knowledgebase in 2026. Nucleic Acids Res. 54, D1779–D1792. 10.1093/nar/gkaf1292

George, B.M., Eleftheriou, M., Yankova, E., Perr, J., Chai, P., Nestola, G., Almahayni, K., Evans, S., Damaskou, A., Hemberger, H., Lebedenko, C.G., Rak, J., Yu, Q., Bapcum, E., Russell, J., Bagri, J., Volk, R.F., Spiekermann, M., Stone, R.M., Giotopoulos, G., Huntly, B.J.P., Baxter, J., Camargo, F., Liu, J., Zaro, B.W., Vassiliou, G.S., Möckl, L., de la Rosa, J., Flynn, R.A., Tzelepis, K., 2026a. Treatment of acute myeloid leukemia models by targeting a cell surface RNA-binding protein. Nat. Biotechnol. 44, 430–443. 10.1038/s41587-025-02648-2

George, B.M., Rodriguez-Pöhnlein, A., DeCastro, D.J., Porat, J., Zhou, B., Abeja, D.M., Lumpkin, R.J., Metivier, R.J., Schmoker, A., Walter, W., Haferlach, T., Pikman, Y., Donovan, K.A., Stegmaier, K., Fischer, E.S., Wachter, F., Flynn, R.A., 2026b. Unbiased, systematic surfaceome profiling of Acute Leukemia to identify novel immunotherapy targets. 10.64898/2026.09.16.752145

Gygi, S.P., Rochon, Y., Franza, B.R., Aebersold, R., 1999. Correlation between protein and mRNA abundance in yeast. Mol. Cell. Biol. 19, 1720–1730. 10.1128/MCB.19.3.1720

Hallgren, J., Tsirigos, K.D., Pedersen, M.D., Armenteros, J.J.A., Marcatili, P., Nielsen, H., Krogh, A., Winther, O., 2022. DeepTMHMM predicts alpha and beta transmembrane proteins using deep neural networks. 10.1101/2022.04.08.487609

Howard, M.K., Hoppe, N., Huang, X.-P., Mitrovic, D., Billesbølle, C.B., Macdonald, C.B., Mehrotra, E., Rockefeller Grimes, P., Trinidad, D.D., Delemotte, L., English, J.G., Coyote-Maestas, W., Manglik, A., 2025. Molecular basis of proton sensing by G protein-coupled receptors. Cell 188, 671–687.e20. 10.1016/j.cell.2024.11.036

Hulo, C., de Castro, E., Masson, P., Bougueleret, L., Bairoch, A., Xenarios, I., Le Mercier, P., 2011. ViralZone: a knowledge resource to understand virus diversity. Nucleic Acids Res. 39, D576–582. 10.1093/nar/gkq901

Inoue, N., Ikawa, M., Isotani, A., Okabe, M., 2005. The immunoglobulin superfamily protein Izumo is required for sperm to fuse with eggs. Nature 434, 234–238. 10.1038/nature03362

Knol, J.C., Lyu, M., Böttger, F., Nunes Monteiro, M., Pham, T.V., Rolfs, F., Vallés-Martí, A., Schelfhorst, T., de Goeij-de Haas, R.R., Bijnsdorp, I.V., Wang, S., Zhang, F., A, J., Westerman, B.A., Sitek, B., Lehtiö, J., Koster, J., IJzermans, J.N.M., van Laarhoven, H.W.M., Bijlsma, M.F., Medema, J.P., Henneman, A.A., Piersma, S.R., Brakenhoff, R.H., Cloos, J., Cordo’, V., de Jong, D., Kazemier, G., Koppers-Lalic, D., Labots, M., Le Large, T.Y.S., Martens, J.W.M., Meijerink, J.P.P., Mumtaz, M., Scheepbouwer, C., Kibbelaar, R.E., Noske, D.P., Steenbergen, R.D.M., van Grieken, N.C.T., van Houdt, W., Giovannetti, E., van Leenders, G.J.L.H., Jonkers, J., Liu, T., Yan, M., Zhan, X., Guo, T., Jimenez, C.R., 2025. The pan-cancer proteome atlas, a mass spectrometry-based landscape for discovering tumor biology, biomarkers, and therapeutic targets. Cancer Cell 43, 1328–1346.e8. 10.1016/j.ccell.2025.05.003

Laurent, A., Allard, A., Fillet, M., 2025. Cell surface proteomics: Analytical challenges and clinical applications in cancer. TrAC Trends Anal. Chem. 184, 118143. 10.1016/j.trac.2025.118143

Leung, K.K., Wilson, G.M., Kirkemo, L.L., Riley, N.M., Coon, J.J., Wells, J.A., 2020. Broad and thematic remodeling of the surfaceome and glycoproteome on isogenic cells transformed with driving proliferative oncogenes. Proc. Natl. Acad. Sci. 117, 7764–7775. 10.1073/pnas.1917947117

Liu, Y., Beyer, A., Aebersold, R., 2016. On the Dependency of Cellular Protein Levels on mRNA Abundance. Cell 165, 535–550. 10.1016/j.cell.2016.03.014

Loes, A.N., Hinman, M.N., Farnsworth, D.R., Miller, A.C., Guillemin, K., Harms, M.J., 2021. Identification and characterization of zebrafish Tlr4 co-receptor Md-2. J. Immunol. Baltim. Md 1950 206, 1046–1057. 10.4049/jimmunol.1901288

Malmström, E., Malmström, L., Hauri, S., Mohanty, T., Scott, A., Karlsson, C., Gueto-Tettay, C., Åhrman, E., Nozohoor, S., Tingstedt, B., Regner, S., Elfving, P., Bjermer, L., Forsvall, A., Doyle, A., Magnusson, M., Hedenfalk, I., Kannisto, P., Brandt, C., Nilsson, E., Dahlin, L.B., Malm, J., Linder, A., Niméus, E., Malmström, J., 2025. Human proteome distribution atlas for tissue-specific plasma proteome dynamics. Cell 188, 2810–2822.e16. 10.1016/j.cell.2025.03.013

McHugh, M.L., 2012. Interrater reliability: the kappa statistic. Biochem. Medica 22, 276–282.

Mighell, T.L., Lehner, B., 2025. A small molecule stabilizer rescues the surface expression of nearly all missense variants in a GPCR. Nat. Struct. Mol. Biol. 32, 2429–2440. 10.1038/s41594-025-01659-6

Muller, M., Gräbnitz, F., Barandun, N., Shen, Y., Wendt, F., Steiner, S.N., Severin, Y., Vetterli, S.U., Mondal, M., Prudent, J.R., Hofmann, R., van Oostrum, M., Sarott, R.C., Nesvizhskii, A.I., Carreira, E.M., Bode, J.W., Snijder, B., Robinson, J.A., Loessner, M.J., Oxenius, A., Wollscheid, B., 2021. Light-mediated discovery of surfaceome nanoscale organization and intercellular receptor interaction networks. Nat. Commun. 12, 7036. 10.1038/s41467-021-27280-x

Piwowar, H., Priem, J., Larivière, V., Alperin, J.P., Matthias, L., Norlander, B., Farley, A., West, J., Haustein, S., 2018. The state of OA: a large-scale analysis of the prevalence and impact of Open Access articles. PeerJ 6, e4375. 10.7717/peerj.4375

Piwowar, H., Priem, J., Orr, R., 2019. The Future of OA: A large-scale analysis projecting Open Access publication and readership. 10.1101/795310

Plasma membrane (HPA) [WWW Document], 2026. URL https://www.proteinatlas.org/humanproteome/subcellular/plas-ma%2Bmembrane (accessed 6.7.26).

Priem, J., Piwowar, H., Orr, R., 2022. OpenAlex: A fully-open index of scholarly works, authors, venues, institutions, and concepts. 10.48550/arXiv.2205.01833

Raybould, M.I.J., Marks, C., Lewis, A.P., Shi, J., Bujotzek, A., Taddese, B., Deane, C.M., 2020. Thera-SAbDab: the Therapeutic Structural Antibody Database. Nucleic Acids Res. 48, D383–D388. 10.1093/nar/gkz827

Rodriguez-Walker, M., Daniotti, J.L., 2017. Human Sialidase Neu3 is S-Acylated and Behaves Like an Integral Membrane Protein. Sci. Rep. 7, 4167. 10.1038/s41598-017-04488-w

Sayers, E.W., Bolton, E.E., Brister, J.R., Canese, K., Chan, J., Comeau, D.C., Connor, R., Funk, K., Kelly, C., Kim, S., Madej, T., Marchler-Bauer, A., Lanczycki, C., Lathrop, S., Lu, Z., Thibaud-Nissen, F., Murphy, T., Phan, L., Skripchenko, Y., Tse, T., Wang, J., Williams, R., Trawick, B.W., Pruitt, K.D., Sherry, S.T., 2022. Database resources of the national center for biotechnology information. Nucleic Acids Res. 50, D20–D26. 10.1093/nar/gkab1112

Schweke, H., Pacesa, M., Levin, T., Goverde, C.A., Kumar, P., Duhoo, Y., Dornfeld, L.J., Dubreuil, B., Georgeon, S., Ovchinnikov, S., Woolfson, D.N., Correia, B.E., Dey, S., Levy, E.D., 2024. An atlas of protein homo-oligomerization across domains of life. Cell 187, 999–1010.e15. 10.1016/j.cell.2024.01.022

Shen, L., Sun, X., Chen, Z., Guo, Y., Shen, Z., Song, Y., Xin, W., Ding, H., Ma, X., Xu, W., Zhou, W., Che, J., Tan, L., Chen, L., Chen, S., Dong, X., Fang, L., Zhu, F., 2023. ADCdb: the database of antibody–drug conjugates. Nucleic Acids Res. 52, D1097–D1109. 10.1093/nar/gkad831

Sim, J., Wright, C.C., 2005. The kappa statistic in reliability studies: use, interpretation, and sample size requirements. Phys. Ther. 85, 257–268.

Tedman, A., Goel, M., Shah, S., Howard, M.K., Chamness, L.M., Bonifasi, A., Adams, I., Gallagher, J.M., Penn, W.D., Nemec, K., McDonald, E.F., Corman, B.N., Robinson, J.P., Post, C.B., Clark, P.L., Babu, M.M., Manglik, A., Kuntz, C.P., Coyote-Maestas, W., Schlebach, J.P., 2026. Efficient experimental characterization of the GPCRome via deep receptor scanning. Nat. Commun. 17, 9796. 10.1038/s41467-026-76564-7

Teixeira, A.A.R., Zhu, H., Kothiwal, D., Cao, R., Mills, A., 2026. OpenAntigens: a structure-aware database for antigen construct design across the human cell-surface and secreted proteome. 10.64898/2026.07.30.741735

Thul, P.J., Åkesson, L., Wiking, M., Mahdessian, D., Geladaki, A., Ait Blal, H., Alm, T., Asplund, A., Björk, L., Breckels, L.M., Bäckström, A., Danielsson, F., Fagerberg, L., Fall, J., Gatto, L., Gnann, C., Hober, S., Hjelmare, M., Johansson, F., Lee, S., Lindskog, C., Mulder, J., Mulvey, C.M., Nilsson, P., Oksvold, P., Rockberg, J., Schutten, R., Schwenk, J.M., Sivertsson, Å., Sjöstedt, E., Skogs, M., Stadler, C., Sullivan, D.P., Tegel, H., Winsnes, C., Zhang, C., Zwahlen, M., Mardinoglu, A., Pontén, F., von Feilitzen, K., Lilley, K.S., Uhlén, M., Lundberg, E., 2017. A subcellular map of the human proteome. Science 356, eaal3321. 10.1126/science.aal3321

UniProt Consortium, 2025. UniProt: the Universal Protein Knowledgebase in 2025. Nucleic Acids Res. 53, D609–D617. 10.1093/nar/gkae1010

Van Auken, K., Schaeffer, M.L., McQuilton, P., Laulederkind, S.J.F., Li, D., Wang, S.-J., Hayman, G.T., Tweedie, S., Arighi, C.N., Done, J., Müller, H.-M., Sternberg, P.W., Mao, Y., Wei, C.-H., Lu, Z., 2014. BC4GO: a full-text corpus for the BioCreative IV GO task. Database J. Biol. Databases Curation 2014, bau074. 10.1093/database/bau074

van Oostrum, M., Müller, M., Klein, F., Bruderer, R., Zhang, H., Pedrioli, P.G.A., Reiter, L., Tsapogas, P., Rolink, A., Wollscheid, B., 2019. Classification of mouse B cell types using surfaceome proteotype maps. Nat. Commun. 10, 5734. 10.1038/s41467-019-13418-5

Vilella, A.J., Severin, J., Ureta-Vidal, A., Heng, L., Durbin, R., Birney, E., 2009. EnsemblCompara GeneTrees: Complete, duplication-aware phylogenetic trees in vertebrates. Genome Res. 19, 327–335. 10.1101/gr.073585.107

Wang, D., Eraslan, B., Wieland, T., Hallström, B., Hopf, T., Zolg, D.P., Zecha, J., Asplund, A., Li, L., Meng, C., Frejno, M., Schmidt, T., Schnatbaum, K., Wilhelm, M., Ponten, F., Uhlen, M., Gagneur, J., Hahne, H., Kuster, B., 2019. A deep proteome and transcriptome abundance atlas of 29 healthy human tissues. Mol. Syst. Biol. 15, e8503. 10.15252/msb.20188503

Wang, Y., Gao, W., Shi, X., Ding, J., Liu, W., He, H., Wang, K., Shao, F., 2017. Chemotherapy drugs induce pyroptosis through caspase-3 cleavage of a gasdermin. Nature 547, 99–103. 10.1038/nature22393

Wei, C.-H., Allot, A., Lai, P.-T., Leaman, R., Tian, S., Luo, L., Jin, Q., Wang, Z., Chen, Q., Lu, Z., 2024. PubTator 3.0: an AI-powered literature resource for unlocking biomedical knowledge. Nucleic Acids Res. 52, W540–W546. 10.1093/nar/gkae235

Westergaard, D., Stærfeldt, H.-H., Tønsberg, C., Jensen, L.J., Brunak, S., 2018. A comprehensive and quantitative comparison of text-mining in 15 million full-text articles versus their corresponding abstracts. PLoS Comput. Biol. 14, e1005962. 10.1371/journal.pcbi.1005962

Wu, T., Jiang, L., Mukhtar, T., Wang, L., Jian, R., Wang, C., Trinh, T., Kriegstein, A.R., Snyder, M., Li, J., 2026. Single-cell proteomic landscape of the developing human brain. Nat. Biotechnol. 44, 1391–1404. 10.1038/s41587-025-02980-7

Xie, Z., Bailey, A., Kuleshov, M.V., Clarke, D.J.B., Evangelista, J.E., Jenkins, S.L., Lachmann, A., Wojciechowicz, M.L., Kropiwnicki, E., Jagodnik, K.M., Jeon, M., Ma’ayan, A., 2021. Gene Set Knowledge Discovery with Enrichr. Curr. Protoc. 1, e90. 10.1002/cpz1.90

Yue, L., Jiang, W., Li, S., Luo, M., Fan, N., Zhan, X., Sun, R., Cheng, H., Xue, Z., Liu, T., Zhou, Q., Chen, K., Lu, T., Guo, F., Li, D., Ge, W., Nie, Z., Lyu, M., A, J., Wang, Y., Chen, Y., Fu, Z., Xiang, N., Li, L., Yu, F., Teo, G.C., Nesvizhskii, A.I., Wang, M., Snyder, M.P., Collins, B.C., Xiao, Q., Aebersold, R., Xu, F., Yang, H., Zhang, S., Han, Y., Zhu, Y., Ji, Y., Li, Y., Guo, T., 2026. Spatial distribution of the proteome in the human body and in cancers. Nature 1–10. 10.1038/s41586-026-10660-y

Zhang, Y., Liu, R., Ni, M., Gill, P., Lee, A.S., 2010. Cell surface relocalization of the endoplasmic reticulum chaperone and unfolded protein response regulator GRP78/BiP. J. Biol. Chem. 285, 15065–15075. 10.1074/jbc.M109.087445

